# Sexually Dimorphic Influence of Circadian Pacemaker Neurons on Behavioral Rhythms

**DOI:** 10.1101/2024.01.31.578273

**Authors:** Aishwarya Ramakrishnan Iyer, Eva Scholz-Carlson, Evardra Bell, Grace Biondi, Shlesha Richhariya, Maria P. Fernandez

## Abstract

The circadian system regulates the timing of multiple molecular, physiological, metabolic, and behavioral phenomena. In *Drosophila,* as in other species, most of the research on how the timekeeping system in the brain controls the timing of behavioral outputs has been conducted in males, or sex has not been included as a biological variable. A critical set of circadian pacemaker neurons in *Drosophila* release the neuropeptide pigment-dispersing factor (PDF), which functions as a key output factor in the network with complex effects on other clock neurons. Lack of *Pdf* or its receptor, *PdfR,* results in most flies displaying arrhythmicity in activity–rest cycles under constant conditions. However, our results show that female circadian rhythms are less affected by mutations in both *Pdf* and *PdfR*. Crispr-Cas9-mediated mutagenesis of *Pdf,* specifically in ventral lateral neurons (LN_v_s), also has a greater effect on male rhythms. We tested the influence of M-cells on the circadian network and showed that speeding up the molecular clock specifically in M-cells led to sexually dimorphic phenotypes, with a more pronounced effect on male rhythmic behavior. Our results suggest that the female circadian system is more resilient to manipulations of M-cells and the PDF pathway, suggesting that circadian timekeeping is more distributed across the clock neuron network in females.

## Introduction

Differences in neuronal circadian timekeeping between sexes remain relatively unexplored, despite the expanding body of research highlighting the influence of sex on the mechanisms underlying neuronal control of behavior [1]. In mammals, steroid hormones display daily, clock-driven changes in abundance, and while these sex hormones are not required to maintain rhythms, they differentially influence the amplitude of activity behavior between the sexes [1, 2]. Furthermore, structural and functional sex differences have been observed in brain areas that receive direct input from the brain’s circadian timekeeping center [2, 3]. Research in humans has also revealed significant sexual dimorphism: men tend to have lower-amplitude endogenous rhythms than women [4], are less resilient to nocturnal sleep disruptions, and spend less time asleep [5].

In mammals, the main circadian pacemaker resides in the suprachiasmatic nuclei (SCN), which in mice consist of a network of ∼20,000 neurons (reviewed in [6]). The *Drosophila* circadian clock network has ∼150 neurons and is the functional equivalent of the mammalian SCN [7, 8]. Each circadian clock neuron has an intracellular molecular timekeeping mechanism based on a transcriptional translational feedback loop: the genes *Clock* (*Clk*) and *cycle* (*cyc*) promote rhythmic transcription of several key genes, including *Period* (*Per*) *and Timeless* (*Tim*), which build up daily and inhibit their own transcription [9]. Multiple kinases that act on components of these clock proteins and can affect the pace of the molecular clock have been identified. One such kinase is *Doubletime* (DBT), which binds to and phosphorylates PER, regulating its nuclear accumulation [10, 11].

The fly clock network consists of *lateral neurons (LNs*), which include ventrolateral (LN_v_), dorsolateral (LN_d_), and lateral posterior neurons (LPNs), as well as three groups of *dorsal neurons* (*DN1, DN2, and DN3*), some of which can be further subdivided [8, 12–15]. The ventral and dorsal LNs are sufficient to produce the normal endogenous bimodal rhythm of sleep and activity [16, 17]. The four small LN_v_s (s-LN_v_s) are usually referred to as morning cells (M-cells) since they control the morning peak of activity under light dark cycles (LDs). These cells are also essential for maintaining rhythmicity under free-running conditions [18, 19]. The evening peak is controlled by the LN_d_s and a *Pdf*-negative LN_v_, the 5^th^ LN_v_ (E-cells) [16, 20, 21]. Some DNs also contribute to the timing and amount of sleep via the modulation of M and E cells [22–25].

The release of the circadian neuropeptide Pigment Dispersing Factor (PDF) by s-LN_v_s is essential for endogenous circadian timekeeping. A *Pdf* null mutation, *Pdf^01^*, results in a substantial fraction of arrhythmic flies [18], desynchronization of molecular oscillations [26, 27], and phase changes in the electrical activity of some clock clusters, most notably the LN_d_s [28]. Loss of *PdfR* also leads to loss of behavioral rhythms [29–31]. Interestingly, PDF and PDFR also regulate behaviors that are sex-specific or sexually dimorphic. Rival-induced long mating durations require PDF expression in s-LN_v_s, PDFR expression in a subset of LN_d_s, and NPF expression in LN_d_s [32]. PDF controls rhythms in the sexually dimorphic pheromone profiles produced by oenocytes [33] and is involved in long-term mating suppression in males [34]. Both PDF and PDFR contribute to geotactic behaviors [29], and the phenotypes of *Pdf*^01^ mutants are sexually dimorphic, with males showing a more extreme negative geotaxis phenotype [35].

Sexual dimorphism in *Drosophila* sleep/wake cycles has been studied mostly under LD cycles. Males exhibit lower levels of activity and more sleep during the light phase [36–38]. This increase in midday sleep is due to the activity of a subset of sleep-promoting DN1s, which are more active in males [36] and receive input from the male-specific P1 neurons that control male courtship [37]. Unlike studies on circadian rhythms, *Drosophila* sleep research often involves only females. Males also have an earlier and more pronounced morning peak and a larger phase angle between the morning peak and the evening peak [39]. Under conditions of constant darkness and temperature (DD), males of several wild-type strains have a small but significant reduction in the free-running period (FRP) relative to females of the same strain [39]. Moreover, males are more likely to retain a bimodal activity pattern in DD [39]. A recent transcriptomic analysis of *fruitless* (*fru*)-expressing neurons revealed clusters that are enriched for circadian clock genes [40]. Interestingly, only the male dataset revealed coexpression of *fru-* and circadian-related genes. A previous study reported that DN1s express the male-specific Fru^M^ protein [41] and that the number of cells in the DN1_p_s cluster is sexually dimorphic [42]. In addition, the E3 subset of LN_d_s has been shown to be dimorphic in its expression of the neuropeptide NPF [43, 44].

Given the sexually dimorphic roles of neuropeptides, including PDF, in other behaviors [45], we asked whether females were similarly affected by manipulations of the *Pdf/PdfR* pathway. We found that female circadian rhythms are less affected by null mutations in both *Pdf* and *PdfR* and that similar effects are observed via CRISPR-Cas9-mediated *Pdf* mutagenesis, specifically in the LN_v_s. Moreover, speeding up the molecular clock in the LN_v_s via expression of *DBT^s^*leads to an advance of the morning peak in males but not in females, and the pace of the free-running period of activity is significantly shortened only in males. Taken together, our results show that the female circadian system is more resilient to manipulations in the PDF pathway and suggest that *Pdf*+ neurons play a more dominant role in the male than in the female circadian network.

## Results

### Mutations in PDF and PDFR lead to sexually dimorphic phenotypes

A null mutation in *Pdf* results in pronounced behavioral phenotypes in *Drosophila* males [18]. We assayed the locomotor activity rhythms of *Pdf^01^*females under free-running conditions (DD) and found that a large proportion of the experimental females were still rhythmic (Fig. 1 A-C). The rhythmic power of experimental flies was significantly reduced in both sexes (Fig. 1D), but the effect was less pronounced in females (Fig. 1E), suggesting that *Pdf^01^* females have more consolidated rhythms than *Pdf^01^* males. Mutant females that were rhythmic had a slightly, but significantly, shorter free-running period than the controls (Table 1). This phenotype was not always observed in experimental males (Table 1), consistent with a recent study [46]. Sleep cycles under DD also appeared to be more consolidated in females (Fig. 1F). *Pdf* mutants have increased sleep, and this effect is mediated by PDF acting on the LN_v_s themselves [47]. We found that both sexes show an increase in total sleep in LD, but the effect was more pronounced in females (Fig. 1G-I). While the increase in sleep in males was most prominent at midday, females exhibited increased sleep throughout most of the light phase (Fig. 1G).

**Figure 1:**
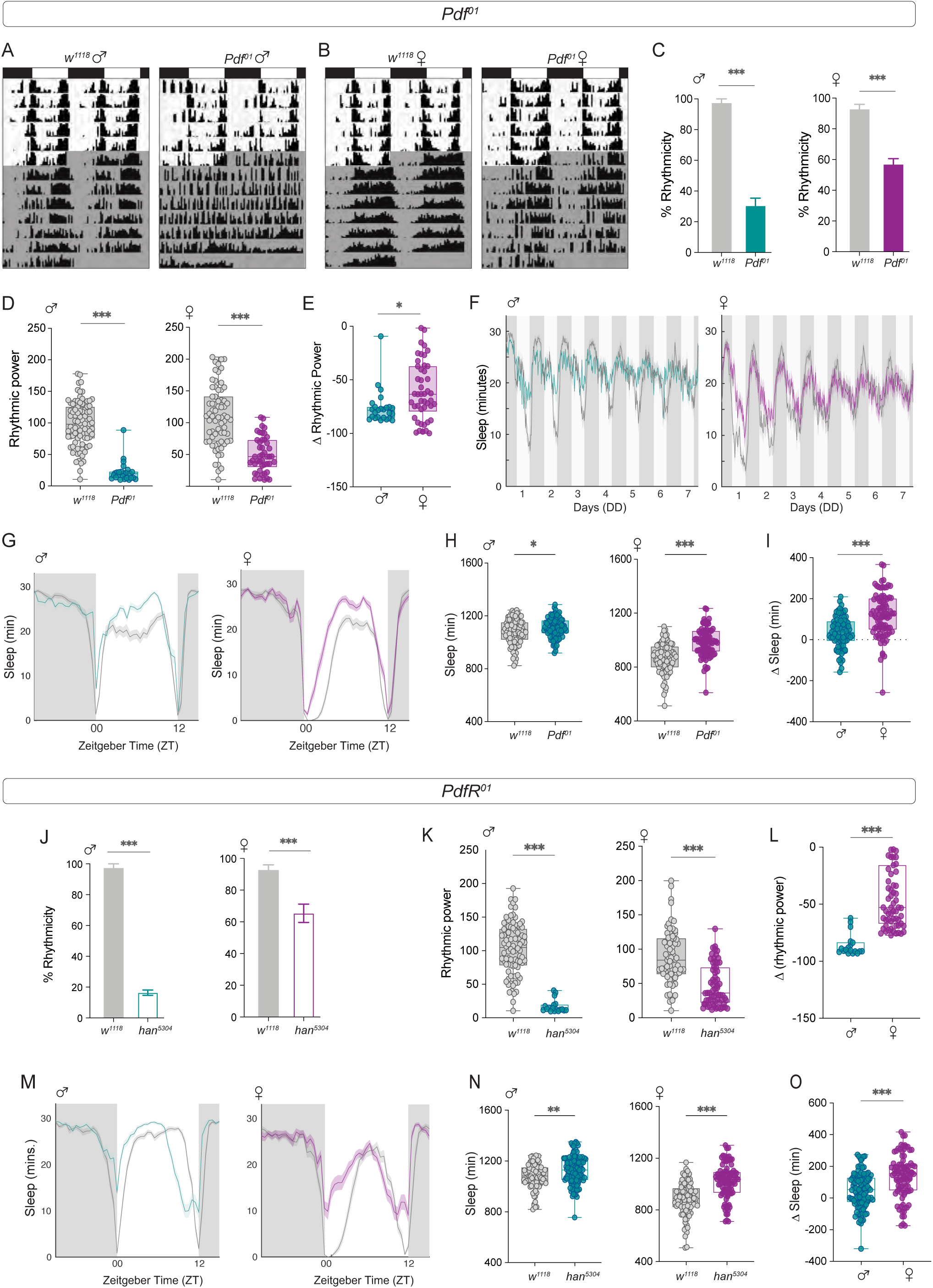
Mutations in *Pdf* and *PdfR* lead to sexually dimorphic phenotypes. **(A)** Representative double-plotted actograms of *w^1118^* (left) and *Pdf^01^*(right) male flies subjected to 6 days of LD followed by seven days of DD. **(B)** Representative actograms of *w^1118^* (left) and *Pdf^01^* (right) female flies subjected to six days of LD followed by seven days of DD. **(C)** Percentages of rhythmic flies are plotted for control (*w^1118^*) and *Pdf^01^* males (left, *n* = 86 (*w^1118^*), *n* = 81 (*Pdf^01^*)) and females (right, *n* = 75 (*w^1118^*), *n* = 101 (*Pdf^01^*)). The error bars represent the SEM values plotted across three replicate experiments. **(D)** Rhythmic power of control (*w^1118^*) and *Pdf^01^* male and females was calculated via a Chi-square Periodogram. **(E)** The differences in rhythmic power between experimental males and females and their respective controls are plotted. **(F)** Average sleep plots of flies over seven days in DD are plotted for male and female control (*w^1118^*) and experimental (*Pdf^01^*) flies. **(G)** Average sleep plots under LD 12:12 for the control (*w^1118^*) and experimental (*Pdf^01^*) groups are plotted for males (left) and females (right). The plots are averaged over flies and days for a period of three days under LD 12:12. **(H)** Total sleep values under LD conditions are plotted for male (left) and female (right) control (*w^1118^*) and experimental (*Pdf^01^*) flies. **(I)** The differences in total LD sleep values between experimental males and females and their respective controls are plotted. **(J)** Percentages of rhythmic flies are plotted for the control (*w^1118^*) and *han^5304^*males (left, *n* = 86 (*w^1118^*), *n* = 115 (*han^5304^*)) and females (right, *n* = 74 (*w^1118^*), *n* = 94 (*han^5304^*)) **(K).** Rhythmic power of the control (*w^1118^*) and *han^5304^* males and females calculated via the chi-square periodogram are plotted. **(L)** The differences in rhythmic power between experimental males and females and their respective controls are plotted. **(M)** Average sleep plots under LD 12:12 for control (*w^1118^*) and experimental (*han^5304^*) flies are plotted for males (left) and females (right). The plots are averaged over flies and days for a period of three days under LD 12:12. **(N)** Total sleep values under LD conditions are plotted for male (left) and female (right) control (*w^1118^*) and experimental (*han^5304^*) flies. **(O)** The differences in total LD sleep values between experimental males and females and their respective controls are plotted. Statistical comparisons were performed between the control and experimental flies of both sexes using a Mann Whitney U test. The box plots extend from the 25^th^ to 75^th^ percentile, with whiskers extending from the smallest to the largest value, and each point represents data from a single fly. Combined data from at least three replicate experiments are plotted. * p < 0.05, ** p < 0.01, *** p < 0.001.

**Table 1:** Table representing the % rhythmicity, phase of the E peak, free-running period and rhythmic power of *w^1118^* and *Pdf^01^* males and females.

| *Pdf<sup>01</sup>* |  |  |  |  |
| --- | --- | --- | --- | --- |
| Genotype | % Rhythmicity ± SEM | E-peak phase ± SEM | Free-running period ± SEM | Rhythmic power ± SEM |
| *w<sup>1118</sup>* (male) | 97.3 ± 2.66 | 11.92 ± 0.07 | 23.8 ± 0.04 | 97.8 ± 3.88 |
| *Pdf<sup>01</sup>* (male) | 30.3 ± 5.05 | 11.72 ± 0.07 | 23.72 ± 0.13 | 22.1 ± 3.24 |
| *w<sup>1118</sup>* (female) | 92.8 ± 3.27 | 11.71 ± 0.07 | 24.1 ± 0.04 | 109.9 ± 5.63 |
| *Pdf<sup>01</sup>* (female) | 56.9 ± 3.63 | 11.85 ± 0.08 | 23.9 ± 0.09 | 49.63 ± 4 |

To rule out the presence of remnant PDF expression in *Pdf^01^* females, we stained the brains of control and experimental males and females with an anti-PDF antibody. We did not observe any traces of PDF in experimental flies of either sex, even with increased laser intensity (Fig. S1A). PDF accumulates rhythmically in the dorsal termini of the s-LN_v_ projections in a time-of-day-dependent manner both in LD and DD [18, 19]. To determine if there were differences in the amplitude of PDF cycling between the sexes in a wild-type background (Canton-S), we dissected control males and females on the third day under DD at 6 timepoints over a 24-hour cycle. Using a COSINOR-based curve fitting method [48], we found that both males and females have clear 24-hour rhythms in PDF cycling in their dorsal projections, with no significant sex differences in amplitude (Fig. S1B-C, Table 2).

**Table 2:**
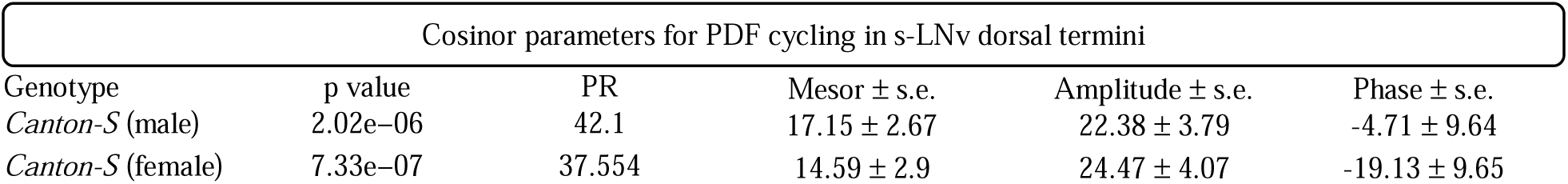
Cosinor analysis parameters for *Canton-S* males and females.

Next, we asked whether the effects of a *Pdf* receptor mutation (*PdfR*) on activity and sleep were also sexually dimorphic. The expression of PDFR, a GPCR, can be detected in most clock neurons, with the exception of 3 LNds, half DN1ps, and some DN3s [49], which coincides with *Cryptochrome* expression in clock neurons [49]. The *han^5304^* mutant is a *PdfR* hypomorph and exhibits *Pdf^01^*-like behavioral phenotypes under both LD 12:12 and DD [29–31]. Under DD, both *Han* males and females showed a significant reduction in rhythmicity compared with the controls, but there was a greater proportion of rhythmic females (∼65%) than males (∼16%) (Fig. 1J). The free-running period of the experimental flies was significantly shorter for both sexes (Table 3), as reported previously for males. Rhythmic power was significantly lower than that of the controls for both *han^5304^* males and females (Fig. 1K), but the effect was more pronounced in males, suggesting that females have more consolidated rhythms (Fig. 1L). Similar to the effect of the *Pdf* mutation, *han^5304^* flies showed significantly higher levels of LD sleep than controls, and this effect was also more pronounced in females (Fig. 1M O). Taken together, these results suggest that female circadian rhythms are less affected by the loss of both *Pdf* and *PdfR*. However, the LD sleep phenotypes were more pronounced in females.

**Table 3:** Table representing the % rhythmicity, phase of the E-peak, free-running period and rhythmic power of *w^1118^* and *PdfR^01^* males and females.

| *PdfR<sup>01</sup>* |  |  |  |  |
| --- | --- | --- | --- | --- |
| Genotype | % Rhythmicity ± SEM | E-peak phase ± SEM | Free-running period ± SEM | Rhythmic power ± SEM |
| *w<sup>1118</sup>* (male) | 97.3 ± 2.66 | 11.68 ± 0.07 | 23.8 ± 0.03 | 103.1 ± 4.15 |
| *PdfR*<sup>01</sup> (male) | 16.4 ± 1.64 | 10.33 ± 0.04 | 21.8 ± 0.12 | 17.46 ± 2.28 |
| *w*<sup>1118</sup> (female) | 92.7 ± 3.25 | 11.45 ± 0.07 | 24.1 ± 0.05 | 89.13 ± 4.75 |
| *PdfR*<sup>01</sup> (female) | 65.4 ± 5.8 | 10.36 ± 0.07 | 22.4 ± 0.07 | 47.3 ± 3.5 |

### CRISPR-Cas9-mediated *Pdf* mutagenesis has more pronounced effects on male behavior

In the *Pdf* null mutant, background effects could contribute to the sexual dimorphism observed in behavioral rhythms. We therefore employed a tissue-specific CRISPR-Cas9-mediated knockout of *Pdf* in both males and females, as described in a recent study that focused on males [50]. To assess the efficiency of the manipulation, we stained for PDF in flies that constitutively expressed *Pdf* gRNA and Cas9 in *Pdf*+ neurons. This experiment was conducted at 28°C, as this temperature was more effective at mimicking the behavioral phenotypes of the *Pdf*^01^ mutant males and allows direct comparison with adult-specific manipulation, which requires switching from 18°C to 28°C.

PDF was reduced in the s-LN_v_s in both sexes (Fig. 2 A-C), although in most experimental brains, we noted faint staining in the dorsal projections of at least one s-LN_v_ in at least one brain hemisphere (Fig. 2A). We quantified PDF intensity in the cell bodies of the sLNvs and found that the signal intensity was reduced in both sexes in a similar manner (Fig. 2B-D). PDF expression within the large LN_v_s was less affected and could be detected in 2–3 l-LNv cell bodies in most brains (Fig. 2A). In addition to behavioral phenotypes, *Pdf*^01^ mutation leads to pronounced misrouting of s-LN_v_ projections in male flies [51] . We employed a transgene expressing a red fluorescent protein under the *Pdf* regulatory sequence [52] and observed faint projections occasionally defasciculating from the main bundle in one or both hemispheres in *Pdf* >*Pdf*-g; *Cas9* flies of both sexes (Fig. 2A, middle panels).

**Figure 2:**
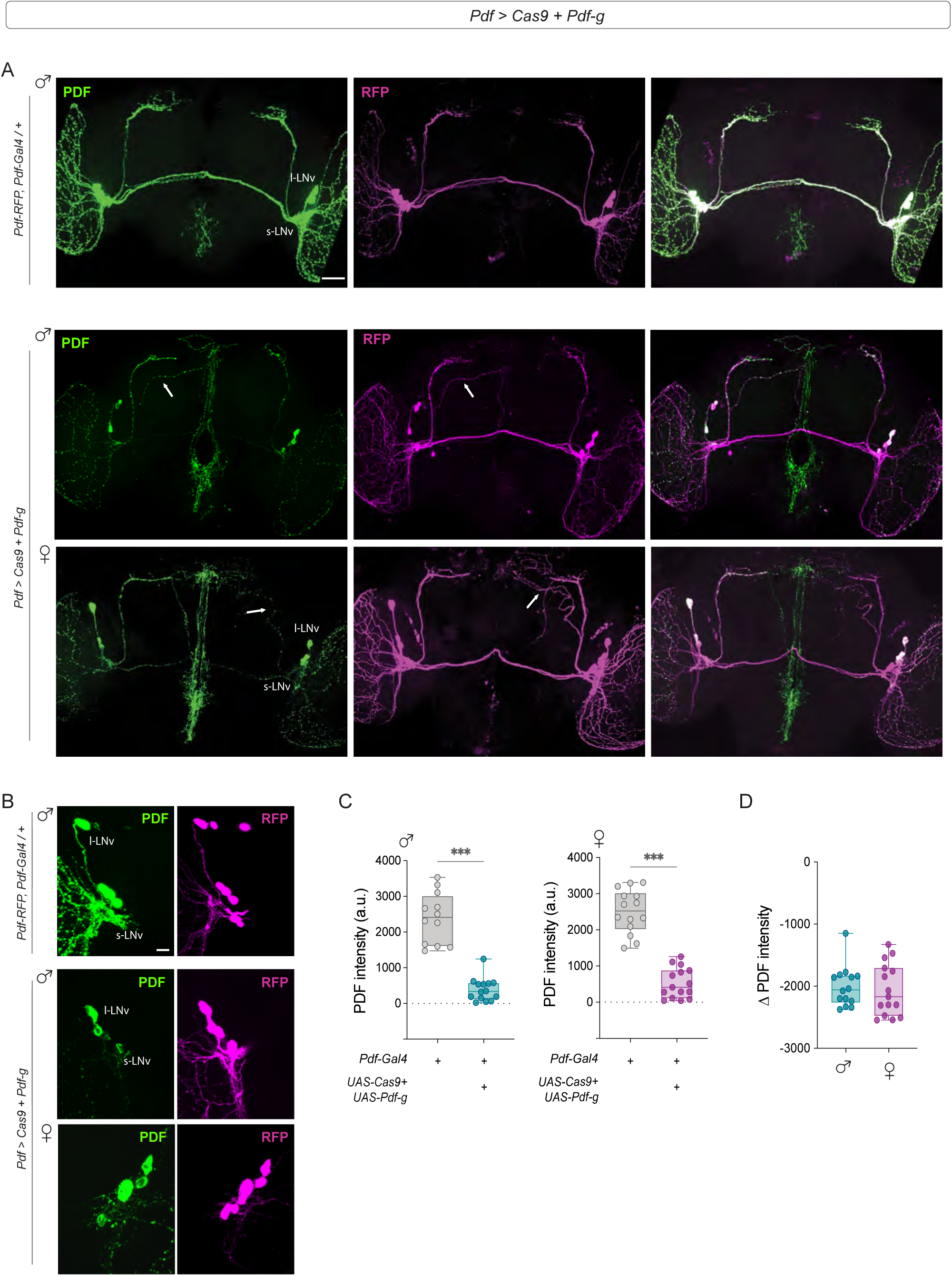
Tissue-specific CRISPR-mediated *Pdf* manipulation leads to a reduction in PDF levels and the misrouting of s-LN_v_ dorsal termini in both sexes. **(A)** Representative confocal images of control (*Pdf-RFP, Pdf-Gal4; tub-Gal80^ts^*) (top) and experimental (*Pdf-RFP, Pdf-Gal4; tub-Gal80^ts^ > Cas9; Pdf-g*) (middle, males; bottom, females) flies stained with RFP and PDF antibodies. Experimental flies show a significant reduction in PDF levels in the sLN_v_s (white arrows, PDF channel) and misrouting of the s-LN_v_ dorsal termini (white arrows, RFP channel). **(B)** Representative confocal images of the small and large LN_v_s of control (*Pdf-RFP, Pdf-Gal4; tub-Gal80^ts^*) (top) and experimental (*Pdf-RFP, Pdf-Gal4; tub-Gal80^ts^ > Cas9; Pdf-g*) (bottom) brains stained with RFP and PDF antibodies. **(C)** Quantification of PDF levels from s-LN_v_ cell bodies in control (*Pdf-RFP, Pdf-Gal4; tub-Gal80^ts^*) and experimental (*Pdf-RFP, Pdf-Gal4; tub-Gal80^ts^ > Cas9; Pdf-g*) flies are plotted for males (left) and females (right). *n* > 12 brains for all genotypes. **(C)** Differences in the PDF intensity values of experimental males and females from their respective parental controls. Statistical comparisons were performed between the control and experimental flies of both sexes using the Mann Whitney U test. The box plots extend from the 25^th^ to 75^th^ percentile, with whiskers extending from the smallest to the largest value, and each point represents data from a single fly. Combined data from at least two replicate experiments are plotted. * p < 0.05, ** p < 0.01, *** p < 0.001. Scale bars = 50 μm.

We analyzed activity rest rhythms in *Pdf* >*Pdf*-g; *Cas9* flies and found that while experimental males were largely arrhythmic, a greater proportion of experimental females remained rhythmic (Fig. 3A-B). The free-running period of the experimental males was not significantly different from that of the controls, but a wide range of periods were observed. Compared with parental controls, *Pdf* >*Pdf*-g; *Cas9* females had significantly shorter free-running periods (Fig. 3C). The rhythmic power was significantly lower in experimental flies of both sexes (Fig. 3D), but the effect was less pronounced in females (Fig. 3E). Sleep under LD was not affected (Fig. S2 A-B), whereas sleep under DD was similarly increased in males and females (Fig. S2 C-E).

**Figure 3:**
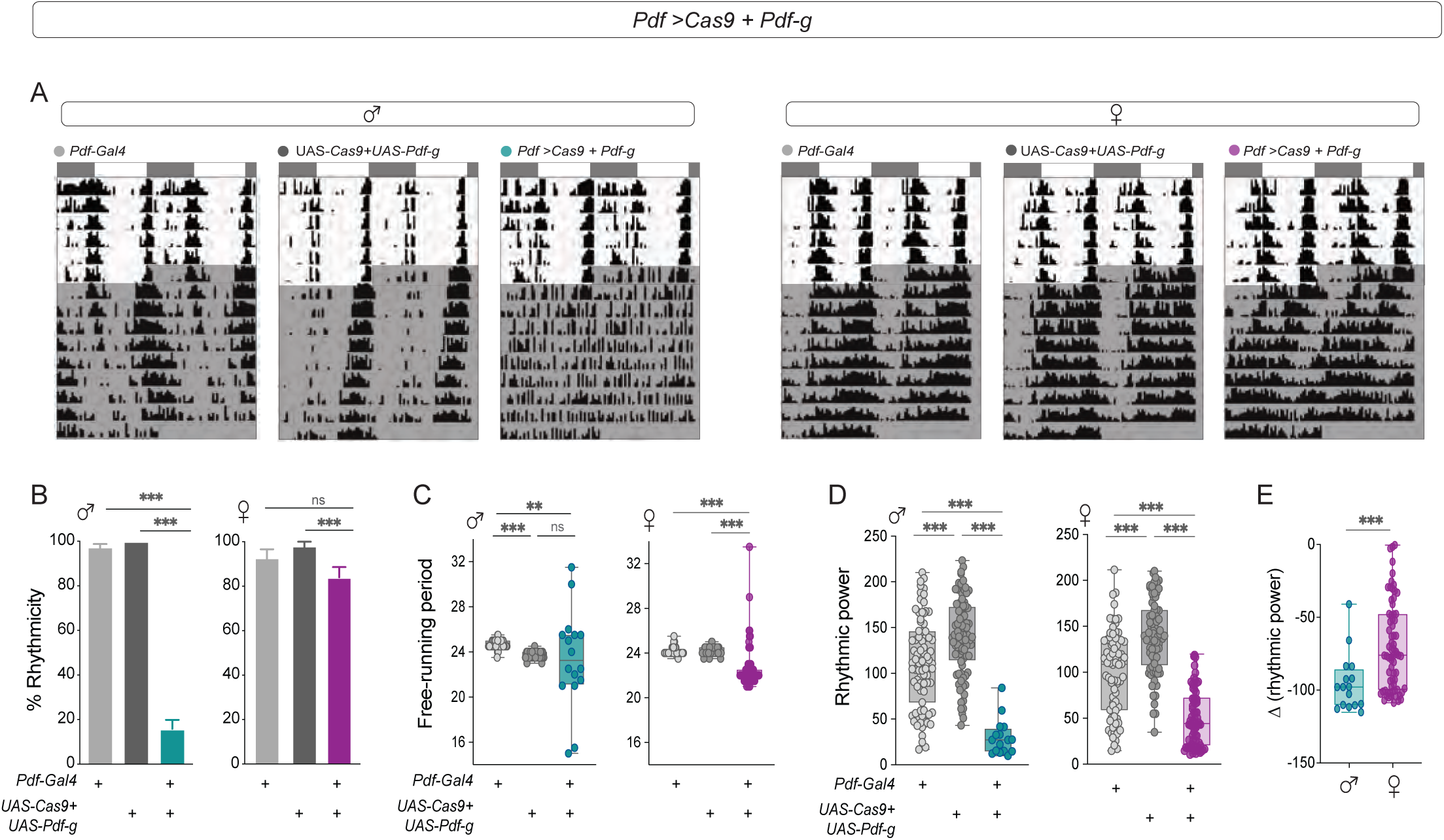
CRISPR-Cas9-mediated *Pdf* mutagenesis has more pronounced effects on male behavior. **(A)** Representative actograms of control (*Pdf-RFP, Pdf-Gal4; tub-Gal80^ts^*) and *(UAS Cas9; Pdf-g)* and experimental (*Pdf-RFP, Pdf-Gal4; tub-Gal80^ts^ > Cas9; Pdf-g*) males (left) and females (right) are plotted for six days of LD followed by nine days of DD. **(B)** Percentages of rhythmic flies are plotted for control (*Pdf-RFP, Pdf-Gal4; tub-Gal80^ts^) and (UAS Cas9; Pdf-g)* and experimental (*Pdf-RFP, Pdf-Gal4; tub-Gal80^ts^ > Cas9; Pdf-g)* males (left, *n* = 85 (*Pdf-RFP, Pdf-Gal4; tub-Gal80^ts^*), *n* = 89 (*UAS Cas9; Pdf-g*), *n* = 106 (*Pdf-RFP, Pdf-Gal4; tub-Gal80^ts^ > Cas9; Pdf-g*) and females (right, *n* = 79 (*Pdf-RFP, Pdf-Gal4; tub-Gal80^ts^*), *n* = 78 (*UAS Cas9; Pdf-g*), *n* = 85 (*Pdf-RFP, Pdf-Gal4; tub-Gal80^ts^ > Cas9; Pdf-g).* The error bars represent SEM values plotted across three replicate experiments. **(C)** Free-running periods of control (*Pdf-RFP, Pdf-Gal4; tub-Gal80^ts^ and UAS Cas9; Pdf-g)* and experimental (*Pdf-RFP, Pdf-Gal4; tub-Gal80^ts^ > Cas9; Pdf-g)* males (left) and females (right) calculated via the chi-square periodogram are plotted. **(D)** Rhythmic power of control (*Pdf-RFP, Pdf-Gal4; tub-Gal80^ts^ and UAS Cas9; Pdf-g)* and experimental (*Pdf-RFP, Pdf-Gal4; tub-Gal80^ts^ > Cas9; Pdf-g)* males (left) and females (right) calculated via the chi-square periodogram are plotted. **(E)** The differences in rhythmic power of experimental males and females from their respective controls are plotted. Flies were kept at 28°C throughout development, and as adults, experiments were conducted at 28°C. Statistical comparisons were performed between the control and experimental flies of both sexes using the Kruskal□Wallis test followed by Dunn’s multiple comparisons test for panels 3B-3D and Mann-Whitney U test for Fig. 3E. The box plots extend from the 25^th^ to 75^th^ percentile, with whiskers extending from the smallest to the largest value, and each point represents data from a single fly. Combined data from at least three replicate experiments are plotted. ** p < 0.01, *** p < 0.001.

We next restricted the CRISPR mutagenesis of *Pdf* to the small LN_v_s via a specific driver from the split-Gal4 collection generated by the Rubin Laboratory (*SS00681*-Gal4). *Pdf* knockdown in the s-LNv resulted in most males being arrhythmic (∼30% rhythmicity), whereas the experimental females were ∼45% rhythmic (Fig. S3A). The percentage of rhythmic flies was significantly lower than that of both controls for experimental males but not for females (Fig. S3A). The free-running period of the experimental males was shorter than that of their respective control flies, whereas the experimental females did not differ from their controls (Fig. S3B). The rhythmic power of the males was lower than that of the controls, but was not different between the experimental females and their controls (Fig. S3C). This suggests that PDF from the s-LNv is important for the behavioral and sex-specific differences observed in *Pdf^01^* mutants.

We next conducted adult-specific CRISPR-Cas9 knockdown of *Pdf* (Fig S4). Although a previous study in males did not establish a connection between developmental misrouting caused by *Pdf* downregulation and the disruption of behavioral rhythms in adulthood [51], sex-specific differences observed upon *Pdf* loss may result from developmental effects. The AGES system was previously employed to rule out the developmental effects of the loss of *Pdf* [50], but a recent study showed that AUXIN exposure induces short- and long-term changes in physiology and behavior [53]. Therefore, we employed a temperature-sensitive Gal80 variant with ubiquitous expression to conditionally inhibit Gal4-mediated expression of the *Pdf* guide and Cas9 [54]. This allows for the temporal control of UAS transgenes since Gal80^ts^ is active at lower temperatures but inactive at higher temperatures. We raised the flies at 18°C and transferred them to 28°C immediately after eclosion so that *Pdf* knockout occurred only in adult flies. Consistent with what was reported for males by Gorostiza et al. [51] via RNA-mediated interference (RNAi) of *Pdf*, we did not observe misrouting in experimental flies (Fig. S4A). While PDF levels were significantly reduced in both males and females, the manipulation was substantially more effective in males (Fig S4B-C). Therefore, although the circadian behavioral phenotypes were again more pronounced in males than in females (Fig S4D-F), this is likely due to higher remaining PDF expression in experimental females.

### Speeding up the M-cell clock leads to a coherent period shortening only in males

Next, we sought to determine if the influence of the *Pdf*-releasing cells themselves was sexually dimorphic. While PDF is released from both large and small LN_v_s, only s-LN_v_s (Morning cells) play key roles in regulating free-running rhythm properties [16]. In males, manipulations that change the pace of the clock specifically in the LN_v_s result in changes in the phase of the morning peak of activity and in the free-running period [20, 55]. We expressed the *Doubletime* ‘short’ (*DBT*^s^) allele [56] under the *Pdf-Gal4* driver (Fig. 4A) and analyzed the effects on behavior in both sexes. We found that *Pdf* > *DBT*^s^ males, but not females, have an advanced phase of the morning peak of activity (M-peak, Fig. 4B-C). This suggests that under LD cycles, the M-oscillator is more effective at setting the phase of male than female behavior.

**Figure 4:**
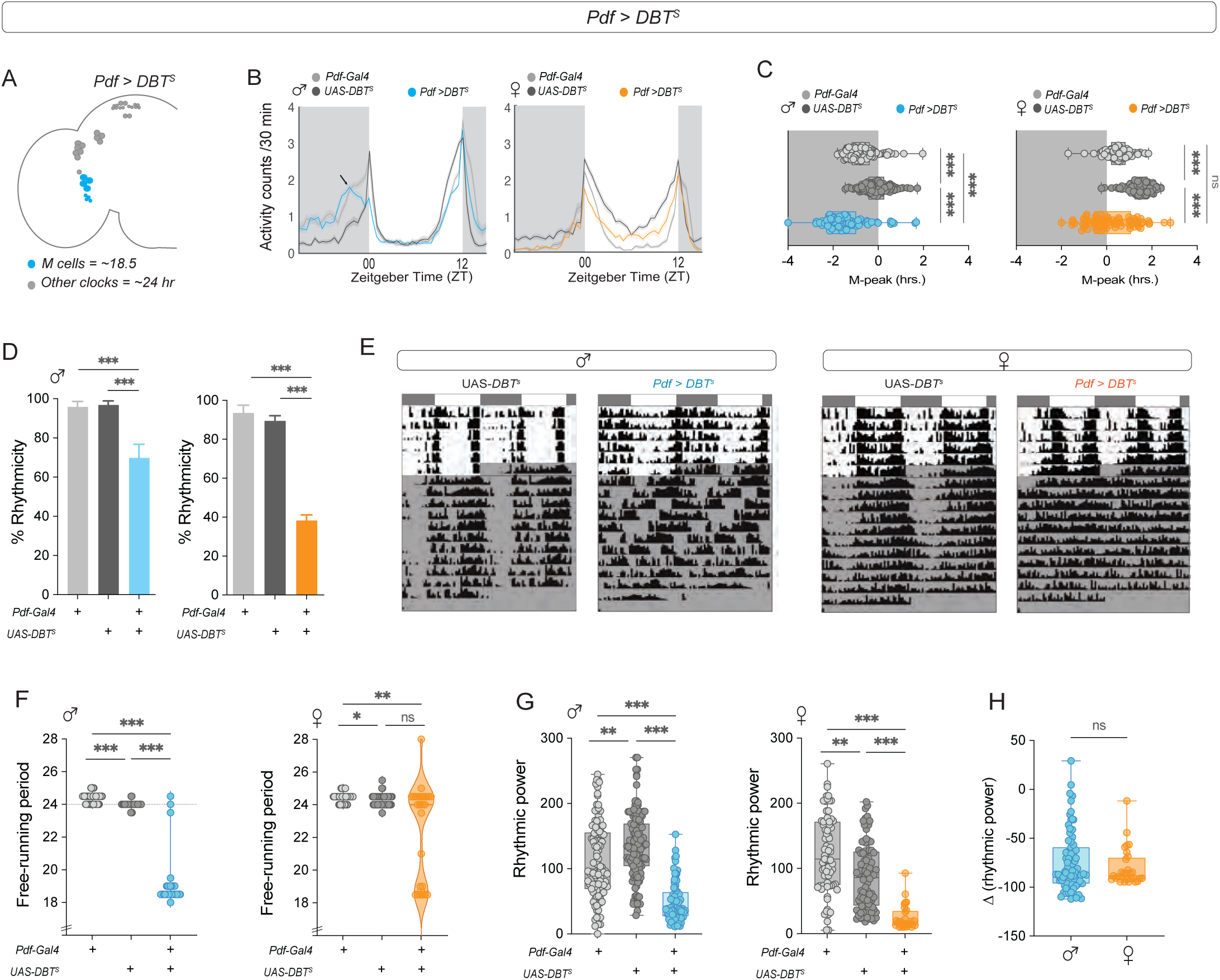
Speeding up the clock in M-cells leads to sexually dimorphic phenotypes. **(A)** Depiction of the adult *Drosophila* brain hemisphere indicating the clock cell subsets (colored) having a faster running molecular clock. **(B)** Average activity plots of control (*Pdf-Gal4*) and (UAS-*DBT^s^*) and experimental (*Pdf > DBT^s^*) flies are plotted for males (left) and females (right). The plots are averaged over flies and days for a period of three days under LD 12:12. **(C)** The phase of the morning peak of activity under LD 12:12 is plotted for control (*Pdf-Gal4*), (UAS-*DBT^s^*) and experimental (*Pdf >DBT^s^*) males (left, *n* = 116 (*Pdf-Gal4*), *n* = 120 (UAS-*DBT^s^*), *n* = 88 (*Pdf > DBT^s^*)) and females (right, *n* = 85 (*Pdf-Gal4*), *n* = 109 (UAS-*DBT^s^*), *n* = 73 (*Pdf > DBT^s^*)). **(D)** Percentages of rhythmic flies are plotted for controls (*Pdf-Gal4), (UAS-DBT^s^),* and experimental (*Pdf > DBT^s^)* males (left, *n* = 90 (*Pdf-Gal4*), *n* = 108 (*UAS-DBT^s^*), *n* = 93 (*Pdf > DBT^s^*) and females (right, *n* = 66 (*Pdf-Gal4*), *n* = 83 (*UAS-DBT^s^*), *n* = 70 (*Pdf > DBT^s^).* The error bars represent the SEM values plotted across three replicate experiments. **(E)** Representative actograms of control (UAS-*DBT^s^*) and experimental (*Pdf > DBT^s^*) males (left) and females (right) plotted for six days of LD followed by ten days of DD. **(F)** Free-running periods of control (*Pdf-Gal4), (UAS-DBT^s^)* and experimental (*Pdf > DBT^s^)* males (left) and females (right) calculated via the chi-square periodogram are plotted. **(G)** Rhythmic power of control (*Pdf-Gal4), (UAS-DBT^s^)* and experimental (*Pdf > DBT^s^)* males (left) and females (right) calculated via the chi-square periodogram are plotted. **(H)** The differences in rhythmic power between experimental males and females and their respective controls are plotted. Statistical comparisons were performed between the control and experimental flies of both sexes using the Kruskal□Wallis test followed by Dunn’s multiple comparisons test for panels 4C, 4F, and 4G and Mann-Whitney U test for Fig. 4H. The box plots extend from the 25^th^ to 75^th^ percentile, with whiskers extending from the smallest to the largest value, and each point represents data from a single fly. Combined data from at least three replicate experiments are plotted. * p < 0.05, ** p < 0.01, *** p < 0.001.

Under free-running conditions, both *Pdf* > *DBT*^s^ male and female flies showed a significantly lower percentage of rhythmic flies than their controls (Fig. 4D), but there were fewer rhythmic females (∼40%) than males (∼65%) (Fig. 4D-E). The free-running period of most *Pdf* > *DBT*^s^ males was ∼18.5 hours (Fig. 4F), whereas the *Pdf* > *DBT*^s^ females showed a large proportion of individuals with a period of ∼24 hours, reflecting the pace of the molecular clock in the rest of the clock network (Fig. 4F). Some *Pdf* > *DBT*^s^ males (∼23%) and a smaller proportion of females (∼6%) also presented complex rhythms (Table 4). The average period value of the second period component (which has a lower power value) was ∼24.07 ± 0.4 for the experimental males and 19.7 ± 0.4 for the experimental females. Neither male nor female control flies exhibited complex rhythms (Table 4). Among rhythmic flies, both *Pdf* > *DBT*^s^ males and females had lower rhythmic power than the controls (Fig. 4G), with no difference between the sexes (Fig. 4H). As a control, we expressed *DBT^s^*via *Clk856-Gal4* which is expressed in most clock neurons (Fig. S5A). Speeding up the molecular clock in most clock neurons significantly advanced both the morning and evening activity peaks of both males and females (Fig. S5B-D). There were no significant differences between experimental males and females in the percentage of rhythmic flies, rhythmic power, or shortening of free-running periods following DBT expression in all clock neurons (Fig. S5E-G).

**Table 4:** Table representing the % rhythmicity, % complex rhythms, free-running period and rhythmic power of *control and Pdf >* males and females.

| *Pdf* > *DBT*<sup>s</sup> |  |  |  |  |
| --- | --- | --- | --- | --- |
| Genotype | % Rhythmicity ± SEM | % Complex rhythms ± SEM | Free-running period ± SEM | Rhythmic power ± SEM |
| *Pdf-Gal4* (male) | 95.86 ± 2.73 | 0 | 24.3 ± 0.03 | 111.8 ± 5.58 |
| *UAS DBT*<sup>s</sup> (male) | 96.79 ± 2.21 | 0 | 23.9 ± 0.01 | 134.8 ± 4.44 |
| *Pdf* > *DBT*<sup>s</sup> (male) | 69.87 ± 6.89 | 23.73 ± 5.51 | 18.9 ± 0.14 | 49.48 ± 3.54 |
| *Pdf-Gal4* (female) | 93.56 ± 4.02 | 0 | 24.4 ± 0.03 | 122 ± 6.76 |
| *UAS DBT*<sup>s</sup> (female) | 89.45 ± 2.74 | 0 | 24.2 ± 0.03 | 89.7 ± 5.19 |
| *Pdf* > *DBT*<sup>s</sup> (female) | 38.41 ± 2.74 | 5.54 ± 3.45 | 22.5 ± 0.58 | 24.8 ± 3.55 |

These results support the notion that M-cells are more dominant in the male than in the female circadian network. Previous studies have shown that blocking synaptic neurotransmission by expressing the tetanus toxin light chain (TeTxLC) in small and large LN_v_s affects male activity rhythms, likely in a *Pdf*-independent manner [57–59]. We analyzed male and female *Pdf* > TeTxLC flies and found that neither sex significantly changed the ability to maintain rhythmicity under free-running conditions (Fig S6A-B). Both sexes significantly lengthened the free-running period, but the effect was more pronounced in males (Fig. S6C-D). Rhythmic power was not affected in experimental flies of either sex (Fig. S6E).

We next asked whether changing the pace of the clock in LNds (E-cells) via *DBT^S^* expression also had sexually dimorphic effects on behavior. These cells can be subdivided into at least 3 different clusters on the basis of their anatomy [8], physiology [55], transcriptomic profiles [13] and connectivity patterns [15]. The PDFR-expressing E1 and E2 clusters have been shown to regulate evening activity under LD [60, 61] and to be able to maintain free-running activity rhythms in the absence of a functional clock in M cells [62, 63], whereas the behavioral role of the E3 cluster remains unknown. We used the MB122-B split-gal4 driver to target the E1 and E2 subsets (Fig. 5A) and found that while expressing *DBT^S^*in this group of evening cells significantly advanced the phase of the E-peak in experimental flies of both sexes (Fig. 5B-C), the effect was more pronounced in females (Fig. 5D). Speeding up of clocks in the E1 + E2 LN_d_s did not significantly alter the free-running period or rhythmic power of experimental flies of either sex (Fig. 5E-F). M-cells have been shown to be the dominant oscillators in DD and to regulate rhythm properties such as persistence and the free-running period of endogenous locomotor rhythms to a large extent [16, 20, 21, 55], although manipulations of other clock cells can affect rhythm properties to some extent [55, 64]. Thus, speeding up the clock in the PDFR^+^ E1 and E2 clusters leads to similar behavioral phenotypes in males and females under free-running conditions. Taken together, these results indicate that the relative influence of the M and E subsets of clock neurons are sexually dimorphic.

**Figure 5:**
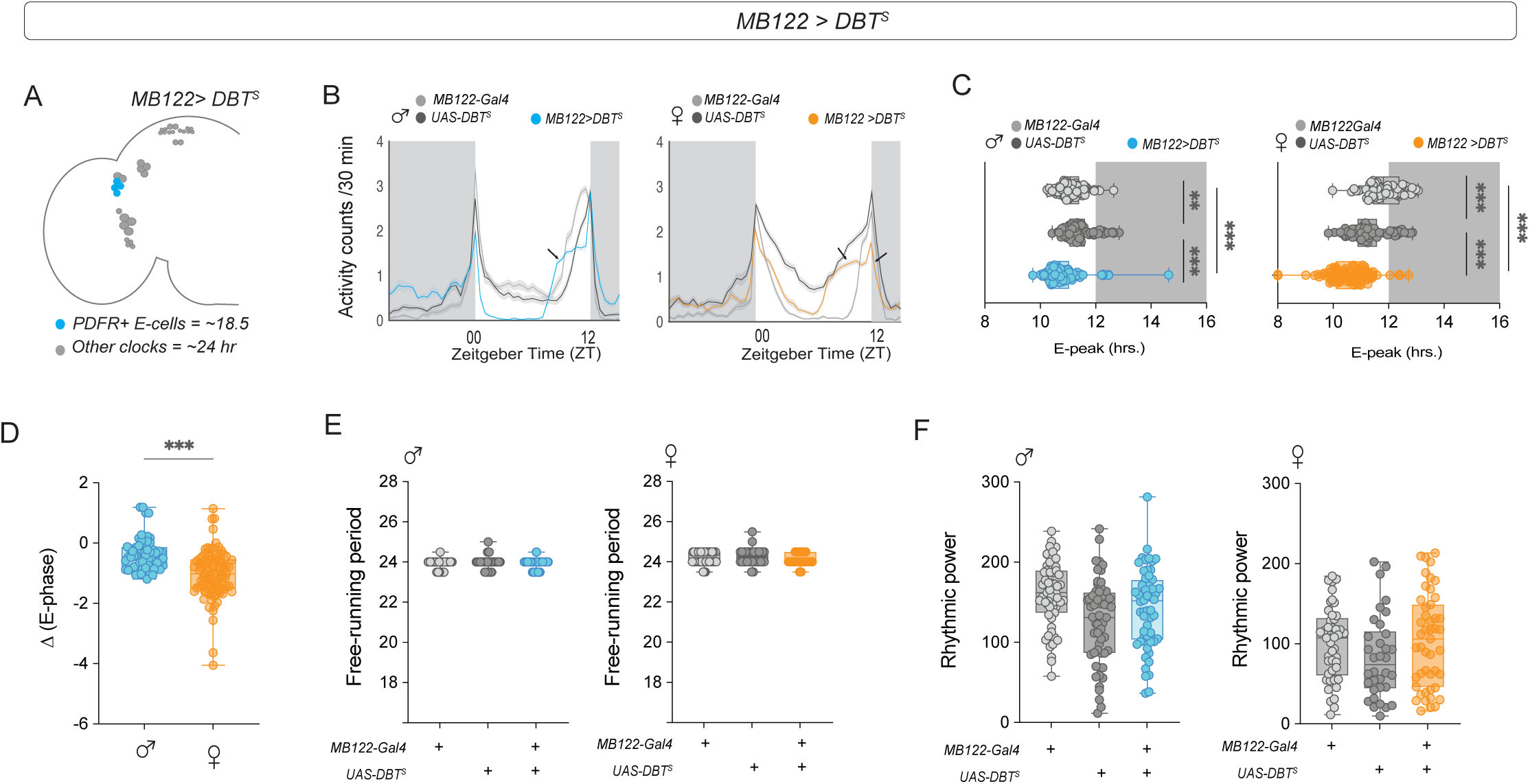
Speeding up the clock in an E-cell subset leads to a more advanced evening peak in females. **(A)** Depiction of the adult *Drosophila* brain hemisphere indicating the clock cell subsets (colored) having a faster running molecular clock. **(B)** Average activity plots of control (*MB122B-Gal4*) and (*UAS-DBT^s^*) and experimental (*MB122B > DBT^s^*) flies are plotted for males (left) and females (right). The plots are averaged over flies and days for a period of three days under LD 12:12. **(C)** Phase of the evening peak of activity under LD 12:12 for controls (*MB122B-Gal4*) and (*UAS-DBT^s^*) and experimental (*MB122B > DBT^s^*) males (left, *n* = 92 (*MB122B-Gal4*), *n* = 93 (*UAS-DBT^s^*), *n* = 93 (*MB122B > DBT^s^*)) and females (right, *n* = 85 (*MB122B-Gal4*), *n* = 87 (*UAS-DBT^s^*), *n* = 89 (*MB122B > DBT^s^*)). **(D)** The differences in the phase of the E peak between experimental males and females and their respective controls are plotted. **(E)** Free-running period of control (*MB122B-Gal4* and *UAS-DBT^s^)* and experimental (*MB122B > DBT^s^)* males (left) and females (right) calculated via the chi-square periodogram are plotted. **(F)** Rhythmic power of control (*MB122B-Gal4 and UAS-DBT^s^)* and experimental (*MB122B > DBT^s^)* males (left) and females (right) calculated via the chi-square periodogram are plotted. Statistical comparisons were performed between the control and experimental flies of both sexes using the Kruskal□Wallis test followed by Dunn’s multiple comparisons test for panels 5C, 5E, and 5F and Mann-Whitney U test for Fig. 5D. The box plots extend from the 25^th^ to 75^th^ percentile, with whiskers extending from the smallest to the largest value, and each point represents data from a single fly. Combined data from at least three replicate experiments are plotted. ** p < 0.01, *** p < 0.001.

## Discussion

The critical importance of considering sex as a biological variable has gained increasing recognition in biomedical research [65, 66]. Bias toward male subjects is particularly prevalent in neuroscience, with single-sex studies using male animals outnumbering those using female animals at a ratio of 5.5:1 [67]. This disparity extends to chronobiology, resulting in a limited understanding of how sex affects circadian organization in the nervous system. However, work from several laboratories has revealed sexual dimorphism within the SCN and in its input and output pathways [1, 68]. Sex differences in SCN morphology have been described in both animal models and humans, and sex differences in SCN electrical activity and steroid hormone receptors have also been reported (reviewed in [2]). Notably, sex differences in the number of SCN neurons that express the neuropeptide vasoactive intestinal polypeptide (VIP) and in *Vip* mRNA *e*xpression have been reported (reviewed in [2]). The roles of the mammalian VIP and the *Drosophila* PDF in circadian physiology are highly similar, although neither these peptides nor their receptors are sequence orthologs [69].

Several studies have shown the importance of PDF in generating coherent rhythms of ∼24-hour periodicity. Here, we report that females lacking *Pdf* or its receptor *PdfR* are more likely to maintain consolidated activity-rest behavior than males. This could be because of sex differences in PDF signaling mechanisms, PDFR expression, or the influence of other clock neurons within the network. We did not detect sex differences in the overall levels or temporal patterns of PDF accumulation, suggesting that these differences might not contribute to the sexually dimorphic effects of the loss of *Pdf* expression. In males, other neuropeptides are known to act in concert with PDF to maintain consolidated rhythms in the network, although none of them have as profound an effect as PDF in regulating activity-rest rhythms in DD [46, 70]. Single mutants of DH31 and CCHamide1 do not affect activity rhythms by themselves, but the double mutants of these neuropeptides along with *Pdf^01^*(*DH31^01^/Pdf^01^* and *Pdf^01^*/*CCHa^SK8^*) are almost completely arrhythmic, suggesting that these neuropeptides act hierarchically in the network, with PDF being at the top of that hierarchy [46, 70]. The importance of PDF relative to other peptides released by clock neurons may also be sexually dimorphic.

To overcome the possible pleiotropic effects of *Pdf^01^*and *PdfR* mutations, we employed a constitutive knockout of *Pdf* in ventral lateral neurons via the CRISPR-Cas9 method. Although CRISPR manipulation was only partially effective at eliminating PDF expression, it nevertheless produced phenotypes reminiscent of those produced by the *Pdf^01^*mutation. We observed faint staining in the dorsal projections of at least one s-LN_v_ in at least one hemisphere in most brains, and PDF staining in a single s-LN_v_ projection reaching the dorsal brain has been shown to be sufficient for behavioral rhythms [71]. Experimental flies in which *Pdf* was knocked out starting at the onset of promoter expression early in development showed extensive misrouting of their dorsal termini, similar to what has been reported for *Pdf^01^* males [51]. Instances of s-LN_v_ misrouting have also been observed in other core clock mutants, such as *per^01^*and *tim^01^* [72] and *cyc^01^* [73]. We observed misrouting in experimental flies of both sexes (data not shown), and no correlation between misrouting and behavioral phenotypes was found by others for *Pdf^01^*males [51]. Importantly, *Pdf > Pdfg;Cas9* manipulation recapitulates the sexually dimorphic circadian phenotypes of *Pdf^01^*mutants: a larger fraction of females are rhythmic, and females exhibit greater rhythm power.

In males, changing the speed of the M cell clock leads to phase changes in the morning peak under LD [60]. To determine whether M-cell manipulations also have sexually dimorphic effects on behavior, we sped up the molecular clock by expressing the *Doubletime short* allele (*DBT^s)^*. Surprisingly, our results revealed that speeding up the clock of M-cells advances the phase of the morning peak in males, whereas the female morning peak phase is not affected. These results support previous studies conducted in males on the role of M-cells in regulating the morning peak of activity [16, 21, 60] and suggest that M-cells are unable to regulate the phase of morning activity in the same way in females. In DD, males had largely coherent short-term rhythms, and the majority (65%) were rhythmic. In contrast, only 40% of the females were rhythmic, and their period showed a bimodal distribution, with most flies exhibiting an ∼24-hour period. These findings further support the notion that M-cells are less dominant than the circadian clock network in females. A possible explanation for this is that other clock neurons are able to “resist” their influence, and the conflict between the fast-paced M cell clock and the ∼24 clock other clock neurons is what leads to greater arrhythmicity in females.

The expression of a tetanus toxin light chain construct in flies blocks neurotransmission by binding and cleaving the synaptic protein Synaptobrevin [74]. Expressing tetanus toxin in ventral lateral neurons does not affect behavioral rhythmicity and lengthens the free-running period in males, as reported in previous studies [58, 59, 75]. The behavioral phenotypes resulting from the blockade of synaptic transmission differ from those resulting from the loss of PDF [18] or the ablation of *Pdf-*expressing neurons [21], possibly because tetanus toxin affects classical transmission and not the dense core vesicle-mediated release of neuropeptides such as PDF [76]. Abrogating the dorsal termini of the small ventral lateral neurons, where most of the output synapses are found [15, 76], does not result in behavioral phenotypes similar to those of *Pdf*^01^ under either LD or DD [22]. Our results show that blocking synaptobrevin-dependent synaptic transmission in M-cells does not affect rhythmicity but rather lengthens the free-running period. The period lengthening is more pronounced in males, supporting the notion that M-cells have a greater influence on the circadian network in males.

*Cryptochrome* and *PdfR*-expressing clusters of evening cells—the sNPF-expressing E1 cluster and the ITP-expressing E2 cluster [77]—have roles in setting the phase of the E-peak under LD and sustaining behavioral rhythms in the absence of a functional molecular clock in M-cells [62, 63]. To test whether these cells have a differential influence on the network in males and females, we expressed *DBT*^S^ under a driver that is expressed specifically in the E1 and E2 subsets of LN_d_s. Our results showed that speeding up the clocks in the E1+E2 clusters resulted in a phase advance in the evening peak of activity in both sexes, but the effect was more pronounced in females. Neither MB122 > *DBT*^S^ males nor females presented phenotypes under free-running conditions. A possible reason for the behavioral differences observed between the sexes could be redundancy in females, such that the network is not as dependent on PDF or M-cells for timekeeping. This finding suggests that the female network could have a more distributed mode of timekeeping throughout the circadian clock network.

Across species, sex differences in the circadian timing system are largely related to the regulation of reproduction-related behaviors. In mammals, the SCN determines the timing of the release of reproductive hormones and influences the timing of mating (reviewed in [68]) and aggression [78]. In *Drosophila*, the circadian clock controls the timing of sex-specific and sexually dimorphic behaviors, such as male courtship [79] and female sexual receptivity [80] and egg laying [81]. This regulation of rhythmic behaviors requires connectivity between clock neurons and downstream sex-specific circuits. For example, the DN1_p_ cluster, which has been shown to be more active in males [36], is functionally connected to the male-specific *fru*-expressing P1 neurons that regulate male courtship [37]. In females, Allostatin C-producing DN1_p_s have been shown to connect to downstream targets to control rhythms in oogenesis [82], and the Janelia female hemibrain connectome revealed that the LN_d_s form connections with the *doublesex*-expressing PC1 cluster [15]. Our data suggest that the relative hierarchy of circadian oscillators is sexually dimorphic, with a less dominant M oscillator in females. This pattern of circadian timekeeping may serve an adaptive purpose, ensuring the precise timing of essential female-specific behaviors crucial for reproductive fitness, such as sexual receptivity and egg laying.

## Acknowledgments

We are grateful to Charlotte Helfrich-Forster, Abhilash Lakshman, Cahir O’Kane, Jeff Price, and Paul Taghert for helpful discussions and Orie Shafer and members of the Fernandez Lab for helpful comments on the manuscript. We also thank Justin Blau, Aljoscha Nern, Gerry Rubin, Michael Rosbash, and Paul Taghert for sharing fly lines. The mouse anti-PDF antibody was obtained from the Developmental Studies Hybridoma Bank, created by the NICHD of the NIH and maintained at The University of Iowa, Department of Biology, Iowa City, IA 52242. Stocks obtained from the Bloomington *Drosophila* Stock Center (NIH P40OD018537) were used in this study. This work was supported by an NIH grant (R01NS118012) and an NSF Grant (IOS-2239994) to M.P.F. and a Beckman Foundation Scholarship to E.S-C.

## Materials and methods

### Fly lines and rearing

All the genotypes were reared on corn syrup soy media (Archon Scientific; Durham, NC) under LD 12:12 cycles at 25^°^C unless specified otherwise (see figure legends for details). The fly lines used in this study were *Canton-S*, *w^1118^*, *Pdf^01^*, *PdfR^01^*, *Pdf-RFP, Pdf-Gal4; tub-Gal80^ts^*, UAS-*Cas9;* UAS*-Pdfg*, *Pdf-Gal4,* UAS*-DBT^s^, UAS TeTxLC, Clk856Gal4, s-LNvGal4,* and *MB122B-Gal4.* See the fly lines and reagents table below for more details.

### Activity□rest behavior recording and analysis

Individual male and virgin female flies (3–5 days old) were housed in glass locomotor tubes containing 2% agar□4% sucrose food on one end and yarn on the other end. Locomotor activity was recorded using Drosophila activity monitors (DAM, Trikinetics, Waltham, United States of America). The experiments were conducted in Tritech or Percival incubators under controlled light and temperature conditions. Flies were entrained to 12:12 LD cycles for at least 5 days and then transferred to constant darkness (DD) for at least 7 days at a constant temperature of 25°C unless otherwise specified (see figure legends for details). The raw data obtained from the DAM system were scanned and binned into activity counts of 15 min intervals via the DAM File scan. The data were analyzed via the CLOCKLAB software (Actimetrics, Wilmette, IL).

The values of period and rhythmic power were calculated for a period of 7 days via a chi-square periodogram with a cutoff of *p =0.01*. The rhythmic power for each designated rhythmic fly was determined by subtracting the chi-square significance value from the power of the periodogram. Flies that did not exhibit a periodicity peak above the significance threshold were categorized as “arrhythmic,” and their period and rhythmic power were not included in the analysis. The values of the morning and evening peaks were calculated via PHASE software [83]. The total LD sleep values for all the genotypes were calculated for a period of 3 days (LD days 2--4) via the PHASE software. Representative actograms were generated via ClockLab, and activity plots were generated via PHASE. The period, rhythmic power, total sleep, and phase values of all the flies for a particular experimental genotype were compared against the background or parental controls via either the Mann□Whitney test or the Kruskal□Wallis ANOVA followed by the Dunn’s multiple comparisons test. The details of the statistical comparisons and the number of flies used in a given experiment are indicated in their respective figure legends. The number of rhythmic flies of the experimental genotype was compared against their respective background or parental controls via Fisher’s exact test. All the statistical analyses were performed via GraphPad Prism 9.0.

### Immunohistochemistry

The brains of adult male or female flies were dissected in ice-cold Schneider’s insect media (S2) and fixed immediately after dissection in 2% paraformaldehyde (PFA) in S2 media for 30 minutes at room temperature. The fixed brains were washed (three washes of 10 minutes each) with 0.3% PBS-Triton X 100 (PBS-TX) and then treated with blocking solution (5% normal goat serum made in 0.3% PBS-TX) for 1 hr at room temperature. The brains were then incubated with primary antibodies at 4°C for 24 hrs. The primary antibodies used were anti-PDF (mouse, 1:3000, C7, DSHB) and anti-RFP (rabbit, 1:2000, Rockland Immunochemicals). After incubation, the brains were subjected to six washes with 0.3% PBS-TX and incubated with Alexa Fluor-conjugated secondary antibodies overnight at 4°C. The following secondary antibodies were used: goat anti-mouse 488 (1:3000, Invitrogen) and goat anti-rabbit 568 (1:3000, Invitrogen). After incubation, the brain samples were washed 6 times with 0.3% PBS-TX, cleaned and mounted on a clean glass slide using Vectashield mounting media.

### Image acquisition and analysis

The brains were imaged via a confocal microscope (Olympus FV3000) with an Olympus UPLanXApo 20x or 40x objective. Image analysis was performed via Fiji software [84]. In the samples, small and large ventral lateral neurons were classified on the basis of their anatomical locations and expression of the PDF. PDF intensities in these cells were measured by selecting the slice of the Z-stack that showed the maximum intensity, drawing a region of interest (ROI) around the cells and measuring their intensities. Three to four separate background values were also measured around each cell, and the final intensity was taken as the difference between the cell intensity and the average background.

For quantification of the PDF in the dorsal projections, a rectangular box was drawn as the ROI starting from the point where the PDF projection turns into the dorsal brain and the intensity is measured. Three to four background values were also measured around the projection. The intensity values obtained from both hemispheres for each cell type for each brain were averaged and used for statistical analysis. PDF intensity from the s-LN_v_ was compared between the experimental and control genotypes via a Mann□Whitney test. To estimate different aspects of rhythmicity in PDF oscillations in the dorsal termini of s-LN_v_ in males and females, we used a COSINOR-based curve-fitting method (Cornelissen, 2014). COSINOR analysis was implemented via the CATCosinor function from the CATkit package written for R [85].

## Fly lines and reagents

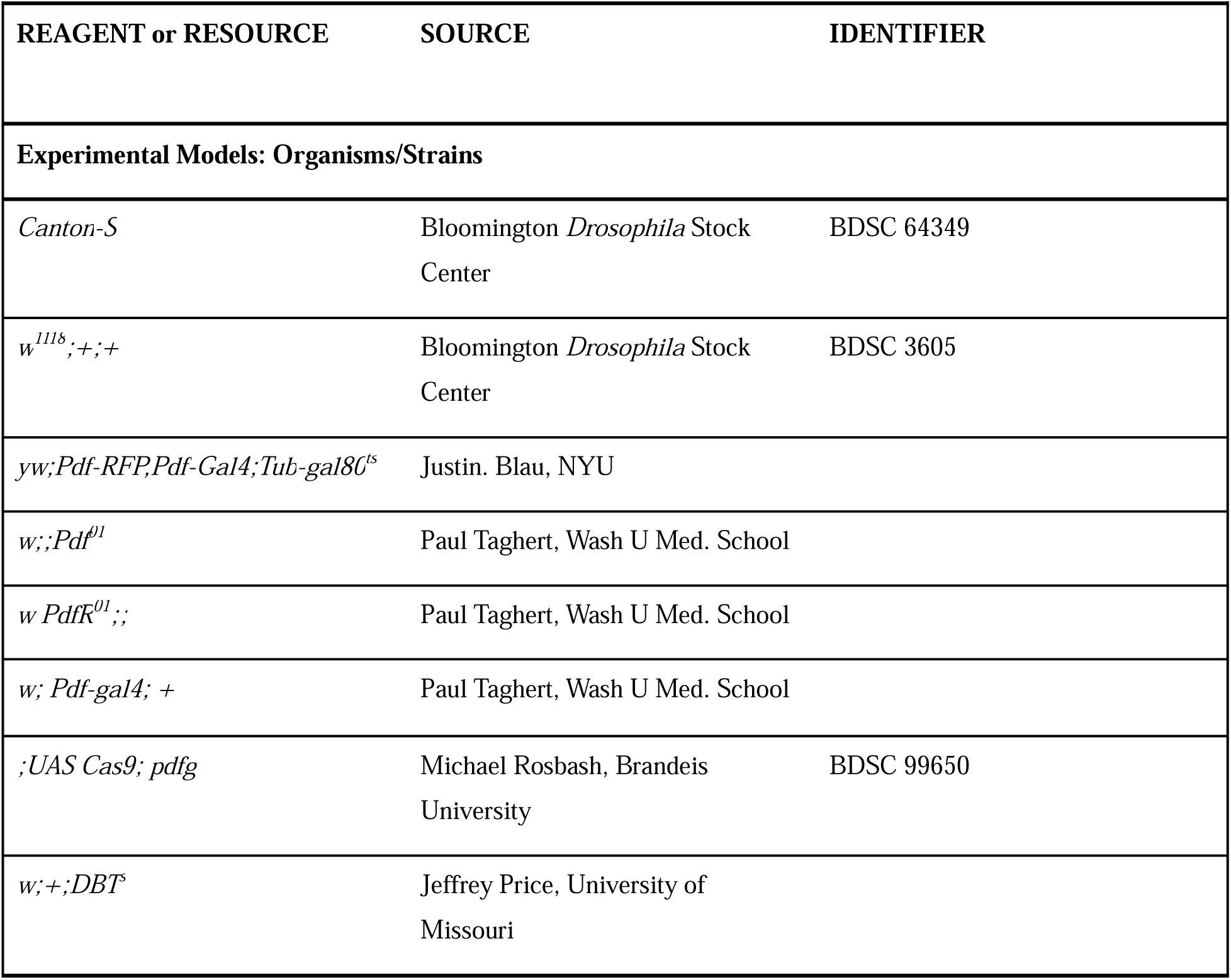
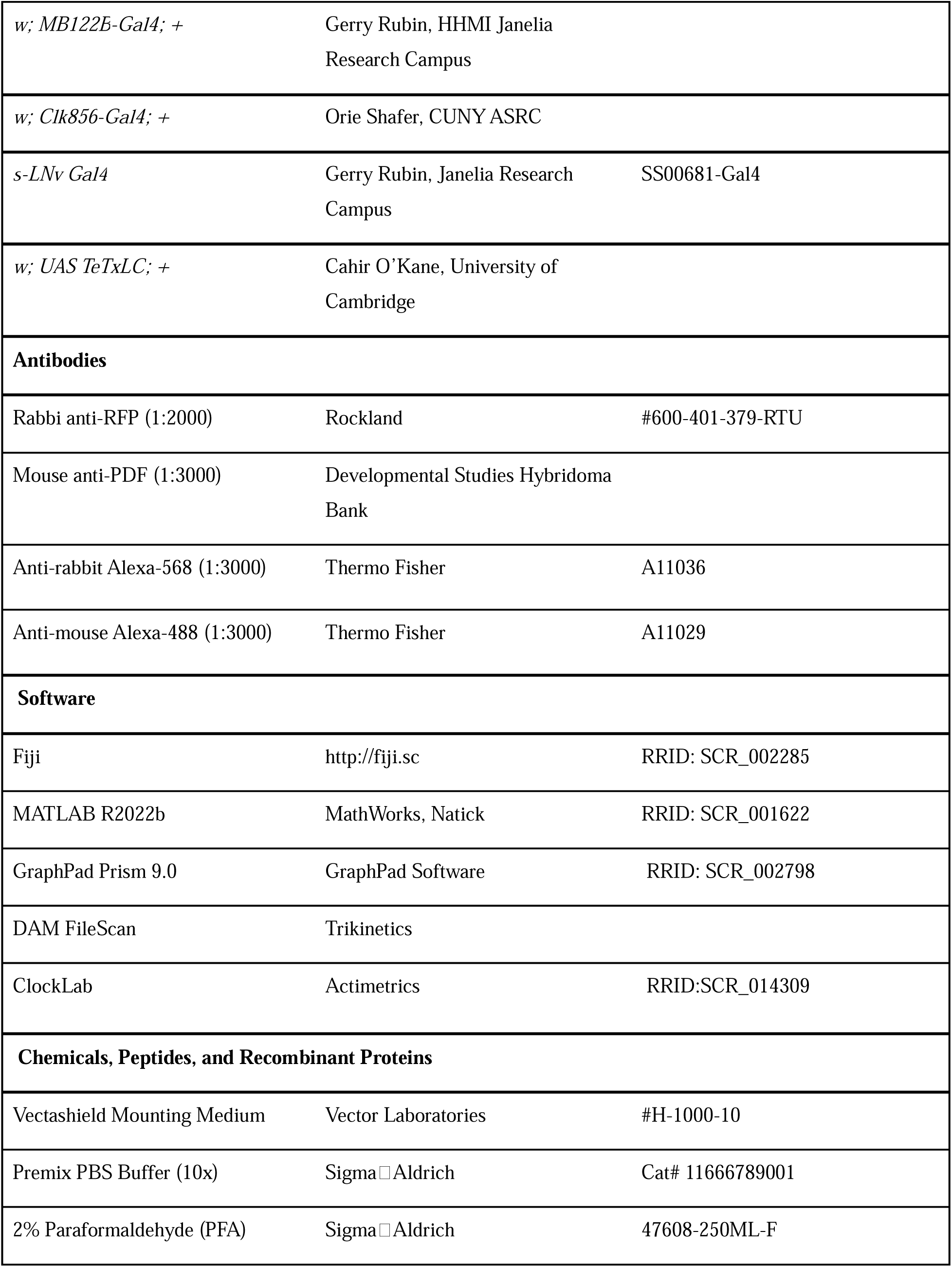
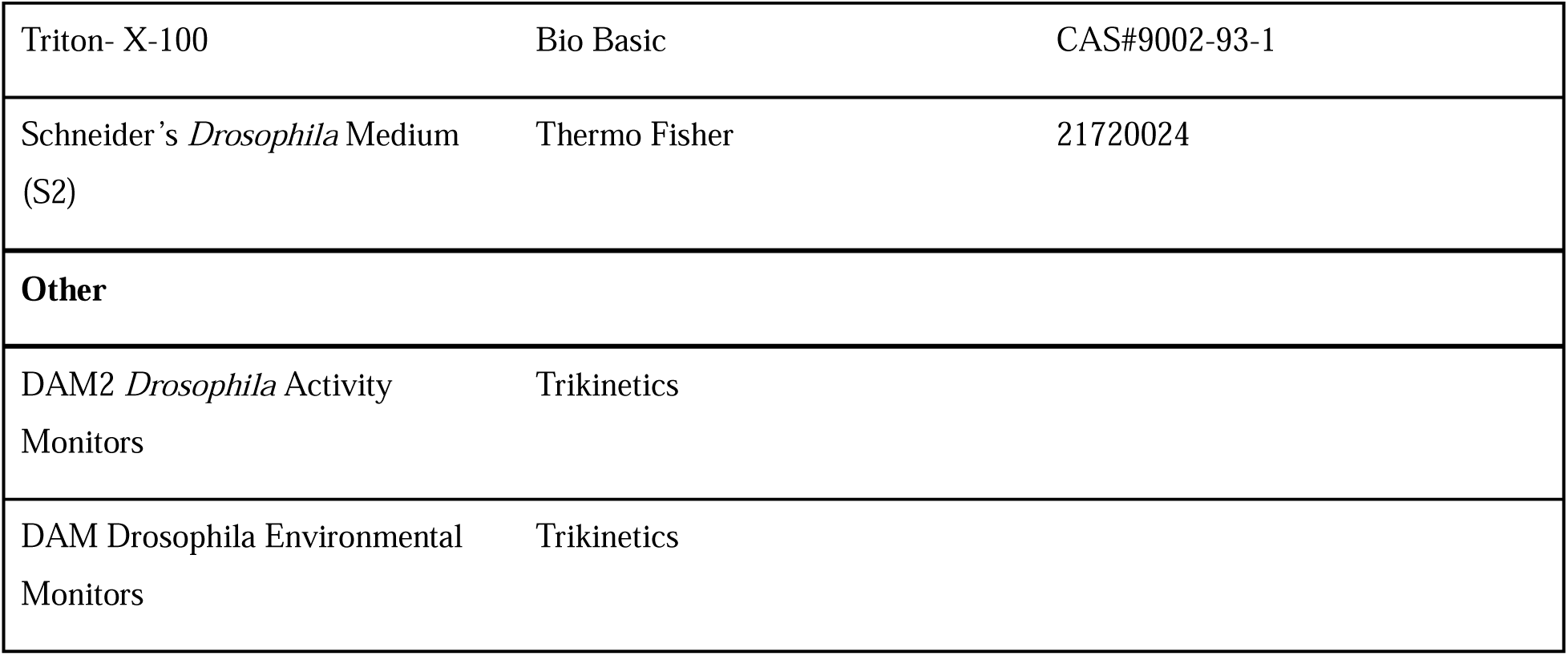

## Supplementary figures

**Figure S1:** PDF rhythmic accumulation is similar in males and females (related to Figure 1.) **(A)** Representative confocal images of control (*w^1118^*) (top) and experimental (*Pdf^01^*) (bottom) male and female flies stained with the PDF antibody. **(B)** Scatterplots of PDF staining intensities of the s-LN_v_ dorsal projections of both male and female flies plotted at different time points over a 24-hr cycle on DD day 3. Each dot represents the mean PDF intensity value averaged over both hemispheres of one brain. The cyan and pink lines are the best fit cosine curves from the parameters that were extracted via COSINOR analysis. See Table 2 for more details. **(C)** Amplitude values of PDF oscillation obtained from COSINOR curve fits are plotted for male and female flies. The error bars are 95% CI values calculated from the standard error obtained from COSINOR analysis. The overlapping error bars indicate that the amplitude values of males and females are not significantly different. n > 7 brain samples/time point.

**Figure S2:** CRISPR-Cas9-mediated *Pdf* mutagenesis has similar effects on male and female sleep. (Related to Figure 3.) **(A)** Average sleep plots under LD 12:12 of control (*Pdf-RFP, Pdf-Gal4; tub-Gal80^ts^ and UAS Cas9; Pdf-g)* and experimental (*Pdf-RFP, Pdf-Gal4; tub-Gal80^ts^ > Cas9; Pdf-g)* males (left) and females (right) are plotted. The plots are averaged over flies and days for a period of 3 days under LD 12:12. **(B)** Total sleep values under LD conditions are plotted for control (*Pdf-RFP, Pdf-Gal4; tub-Gal80^ts^ and UAS Cas9; Pdf-g)* and experimental (*Pdf-RFP, Pdf-Gal4; tub-Gal80^ts^ > Cas9; Pdf-g)* males (left) and females (right). **(C)** Average sleep plots of flies over seven days in DD are plotted for male and female control (*Pdf-RFP, Pdf-Gal4; tub-Gal80^ts^ and UAS Cas9; Pdf-g*) and experimental (*Pdf-RFP, Pdf-Gal4; tub-Gal80^ts^ > Cas9; Pdf-g*) flies. **(D)** Total sleep values under DD conditions are plotted for control (*Pdf-RFP, Pdf-Gal4; tub-Gal80^ts^ and UAS Cas9; Pdf-g)* and experimental (*Pdf-RFP, Pdf-Gal4; tub-Gal80^ts^* > *Cas9; Pdf-g)* males (left) and females (right). **(E)** The differences in the total DD sleep values of experimental males and females from their respective controls are plotted. Statistical comparisons were performed between the control and experimental flies of both sexes using the Kruskal□Wallis test followed by Dunn’s multiple comparisons test for panels S2B and S2D and Mann-Whitney U test for Panel S2E. The box plots extend from the 25^th^ to 75^th^ percentile, with whiskers extending from the smallest to the largest value, and each point represents data from a single fly. Combined data from at least three replicate experiments are plotted. ** p < 0.01, *** p < 0.001.

**Figure S3:** CRISPR-Cas9-mediated *Pdf* mutagenesis in small ventral lateral neurons has more pronounced effects on male circadian behavior. (A) Percentages of rhythmic flies are plotted for control (*s-LNv-Gal4) and (UAS Cas9; Pdf--g)* and experimental (*s-LNv > Cas9; Pdf--g)* males (left, *n* = 16 for all genotypes) and females (right, *n* = 16 for all genotypes). **(B)** Free-running period of control (*s-LNv-Gal4 and UAS Cas9; Pdf-g)* and experimental (*s-LNv > Cas9; Pdf-g)* males (left) and females (right) calculated via the chi-square periodogram are plotted. **(C)** Rhythmic power of control (*s-LNv-Gal4 and UAS Cas9; Pdf-g)* and experimental (*s-LNv > Cas9; Pdf-g)* males (left) and females (right) calculated via the chi-square periodogram are plotted. * p < 0.05, ** p < 0.01, *** p < 0.001. Statistical comparisons were performed between the control and experimental flies of both sexes using the Kruskal□Wallis test followed by Dunn’s multiple comparisons test. The box plots extend from the 25^th^ to 75^th^ percentile, with whiskers extending from the smallest to the largest value, and each point represents data from a single fly. * p < 0.05, ** p < 0.01, *** p < 0.001.

**Figure S4:** Adult-specific CRISPRDCas9-mediated *Pdf* mutagenesis is more effective in males. (Related to Figure 3.) **(A)** Representative confocal images of control (*Pdf-RFP, Pdf-Gal4; tub-Gal80^ts^*) (top) and experimental (*Pdf-RFP, Pdf-Gal4; tub-Gal80^ts^ > Cas9; Pdf-g*) male (middle) and female (bottom) flies stained with RFP and PDF antibodies. The experimental flies presented a significant reduction in PDF levels but no misrouting of the s-LN_v_ dorsal termini. **(B)** Quantification of PDF intensity from s-LN_v_ cell bodies in control (*Pdf-RFP, Pdf-Gal4; tub-Gal80^ts^*) and experimental (*Pdf-RFP, Pdf-Gal4; tub-Gal80^ts^ > Cas9; Pdf-g*) flies are plotted for males (left) and females (right). The data points represent the intensity from a single brain sample. *n* > 10 brains for all genotypes. **(C)** The differences in PDF intensity values of experimental males and females from their respective controls are plotted. **(D)** Percentages of rhythmic flies are plotted for control (*Pdf-RFP, Pdf-Gal4; tub-Gal80^ts^)* and *(UAS Cas9; Pdf-g)* and experimental (*Pdf-RFP, Pdf-Gal4; tub-Gal80^ts^ > Cas9; Pdf-g)* males (left, *n* = 103 (*Pdf-RFP, Pdf-Gal4; tub-Gal80^ts^*), *n* = 109 (*UAS Cas9; Pdf-g*), *n* = 115 (*Pdf-RFP, Pdf-Gal4; tub-Gal80^ts^ > Cas9; Pdf-g*) and females (right, *n* = 96 (*Pdf-RFP, Pdf-Gal4; tub-Gal80^ts^*), *n* = 90 (*UAS Cas9; Pdf-g*), *n* = 119 (*Pdf-RFP, Pdf-Gal4; tub-Gal80^ts^ > Cas9; Pdf-g).* **(E)** Free-running periods of control (*Pdf-RFP, Pdf-Gal4; tub-Gal80^ts^ and UAS Cas9; Pdf-g)* and experimental (*Pdf-RFP, Pdf-Gal4; tub-Gal80^ts^ > Cas9; Pdf-g)* males (left) and females (right) calculated via the chi-square periodogram are plotted. **(F)** Rhythmic power of control (*Pdf-RFP, Pdf-Gal4; tub-Gal80^ts^ and UAS Cas9; Pdf-g)* and experimental (*Pdf-RFP, Pdf-Gal4; tub-Gal80^ts^ > Cas9; Pdf-g)* males (left) and females (right) calculated via the chi-square periodogram are plotted. Flies were raised at 18°C until eclosion and were then transferred to 28°C as adults. The flies were kept at 28°C for a period of four to five days before their activity-rest behavior was recorded at 28°C. Statistical comparisons were performed between the control and experimental flies of both sexes using the Kruskal□Wallis test followed by Dunn’s multiple comparisons test for panels S4E-F and Mann-Whitney U test for panels S4B-C. The box plots extend from the 25^th^ to 75^th^ percentile, with whiskers extending from the smallest to the largest value, and each point represents data from a single fly. Combined data from at least three replicate experiments are plotted. Scale bars represent 50 μm. * p < 0.05, ** p < 0.01, *** p < 0.001.

**Figure S5:** Speeding up the clock in all clock cells leads to a phase advance and shortening of the free-running period of activity rhythms (related to Figures 4 and 5). **(A)** Diagram of an adult *Drosophila* brain hemisphere indicating the clock cell subsets (blue) expressing DBT^s^. **(B)** Average activity plots of control (*Clk856-Gal4*) and (UAS-*DBT^s^*) and experimental (*Clk856 > DBT^s^)* flies are plotted for males (left) and females (right). The plots are averaged over flies and days for a period of three days under LD 12:12. **(C)** Phases of the morning peak of activity under LD 12:12 for controls (*Clk856-Gal4*) and (*UAS-DBT^s^*) and experimental (*Clk856 > DBT^s^*) males (left, *n* = 62 (*Clk856-Gal4*), *n* = 57 (*UAS--DBT^s^*), *n* = 60 (*Clk856 > DBT^s^*)) and females (right, *n* = 59 (*Clk856-Gal4*), *n* = 59 (*UAS--DBT^s^*), *n* = 50 (*Clk856 > DBT^s^*)) are plotted. **(D)** Phase of the evening peak of activity under LD 12:12 for controls (*Clk856--Gal4*) and (*UAS-DBT^s^*) and experimental (*Clk856 > DBT^s^*) males (left) and females (right) are plotted. **(E)** Percentages of rhythmic flies are plotted for control (*Clk856--Gal4*) and (*UAS-DBT^s^*) and experimental (*Clk856 > DBT^s^*) males (left) and females (right). The error bars represent the SEM values plotted across two replicate experiments. **(F)** Free-running periods of rhythmic flies calculated via the chi-square period are plotted for controls (*Clk856--Gal4*) and (*UAS-DBT^s^*) and experimental (*Clk856 > DBT^s^*) males (left) and females (right). **(G)** Rhythmic power of flies calculated via the chi-square periodogram is plotted for controls (*Clk856--Gal4*) and (*UAS-DBT^s^*) and experimental (*Clk856 > DBT^s^*) males (left) and females (right). Statistical comparisons were performed between the control and experimental flies of both sexes via Kruskal□Wallis test followed by Dunn’s multiple comparisons test. The box plots extend from the 25^th^ to 75^th^ percentile, with whiskers extending from the smallest to the largest value, and each point represents data from a single fly. Combined data from at least two replicate experiments are plotted. * p < 0.05, ** p < 0.01, *** p < 0.001.

**Figure S6:** Blocking neurotransmission in *Pdf*-expressing cells leads to more pronounced lengthening of the free-running period in males. (Related to Figure 6). **(A)** Representative actograms of control *(Pdf-Gal4*), and (*UAS TeTxLC*) and experimental (*Pdf > TeTxLC*) males (left) and females (right) are plotted for five days of LD followed by eight days of DD. **(B)** Percentage of rhythmic flies are plotted for control (*Pdf-Gal4*), and (*UAS TeTxLC*), and experimental (*Pdf > TeTxLC*) males (left, n = 56 (*Pdf-Gal4*), n = 51 (UAS *TeTxLC*), n = 52 (*Pdf > TeTxLC*) and females (right, n = 53 (*Pdf-Gal4*), n = 52 (*UAS TeTxLC*), n = 52 (*Pdf > TeTxLC*). Error bars are SEM values plotted across two replicate experiments. **(C)** Free-running period of control (*Pdf-Gal4* and UAS *TeTxLC*) and experimental (*Pdf > TeTxLC*) males (left) and females (right) calculated using the Chi-square Periodogram are plotted. **(D)** The difference in free-running period of experimental males and females from their respective controls are plotted. **(E)** Rhythmic power of control (*Pdf-Gal4* and *UAS TeTxLC*) and experimental (*Pdf > TeTxLC*) males (left) and females (right) calculated using the Chi-square periodogram are plotted. Statistical comparisons were performed between the control and experimental flies for both sexes using the Kruskal-Wallis test followed by the Dunn’s multiple comparisons test for all panels except S6D where comparisons were made using the Mann Whitney U test. The box plots extend from the 25^th^ to 75^th^ percentile, with whiskers extending from the smallest to the largest value, and each point representing data from a single fly. Combined data from at least two replicate experiments are plotted.

## Notes

### Competing Interest Statement

The authors have declared no competing interest.

### Summary of Updates

Affiliations of the first, second, and corresponding author changed from Barnard College to Indiana University. A table was missing in the previous version. Some experiments have been added as supplementary figures. The title has been updated.

