## Supplementary figures and images for "Sexually Dimorphic Influence of Circadian Pacemaker Neurons on Behavioral Rhythms"

### Supplementary Figures

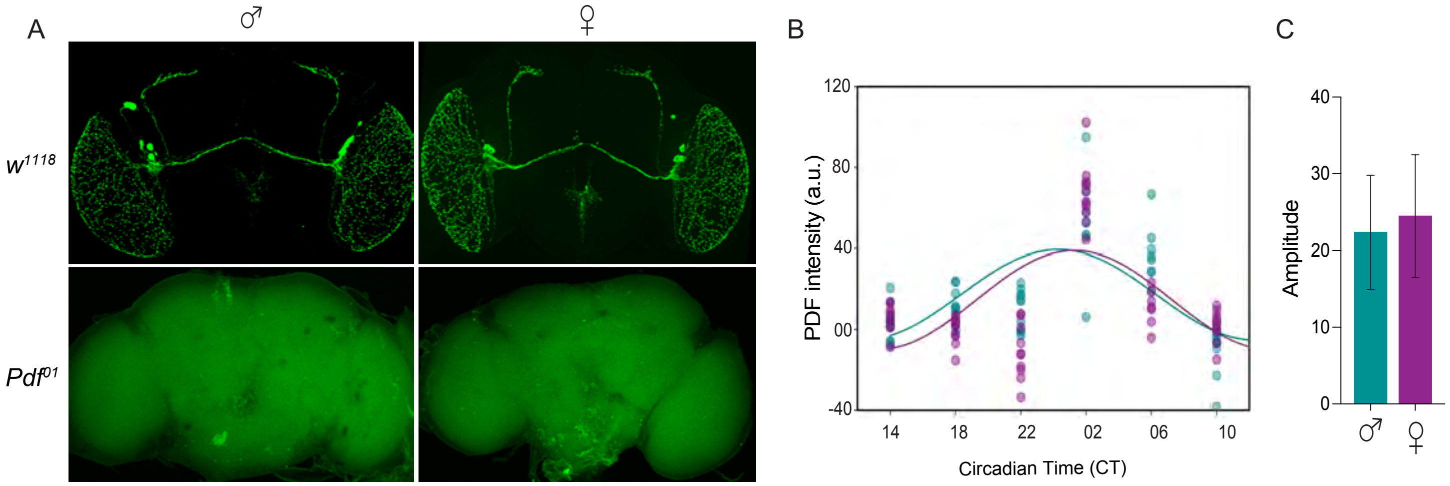

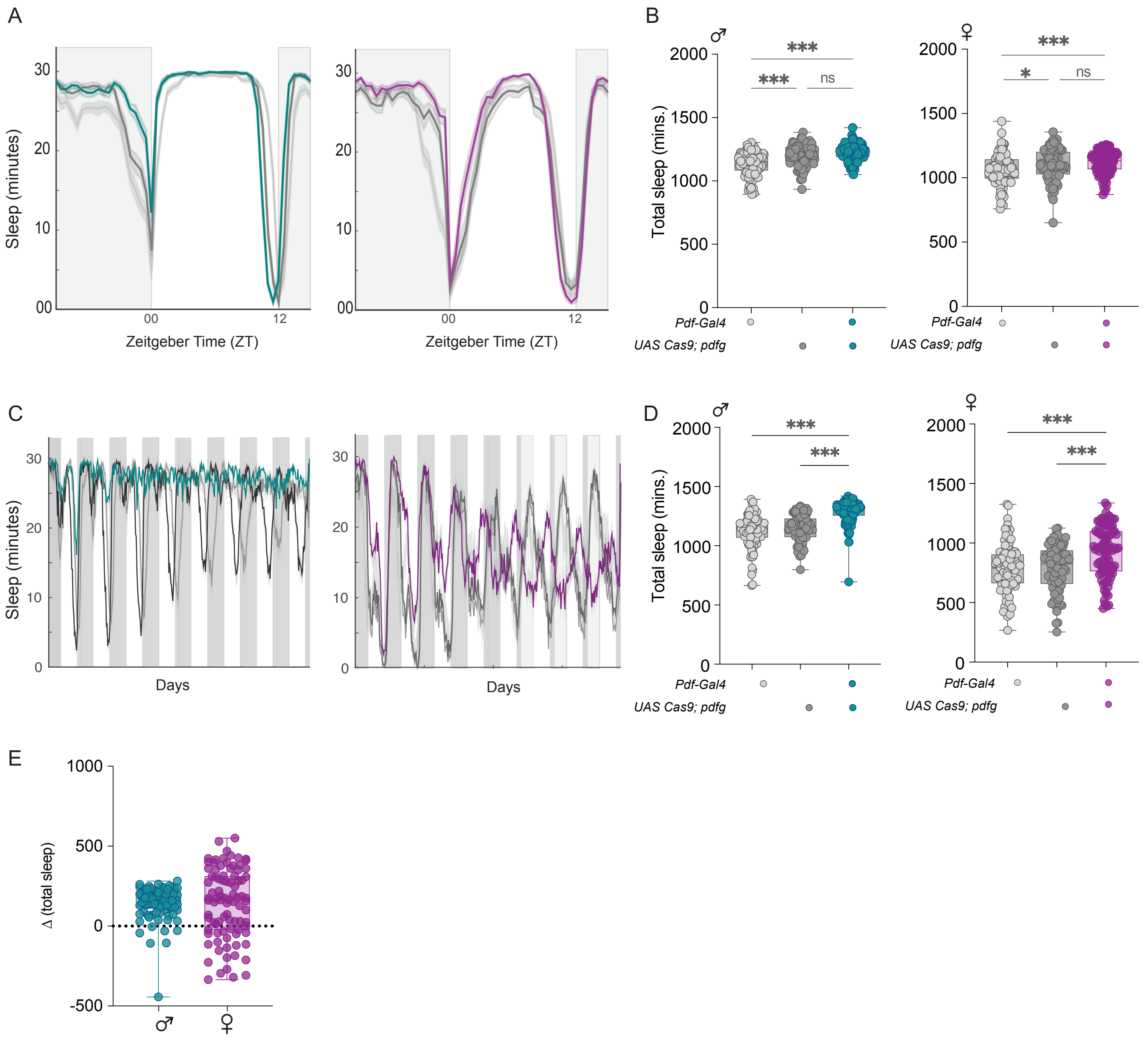

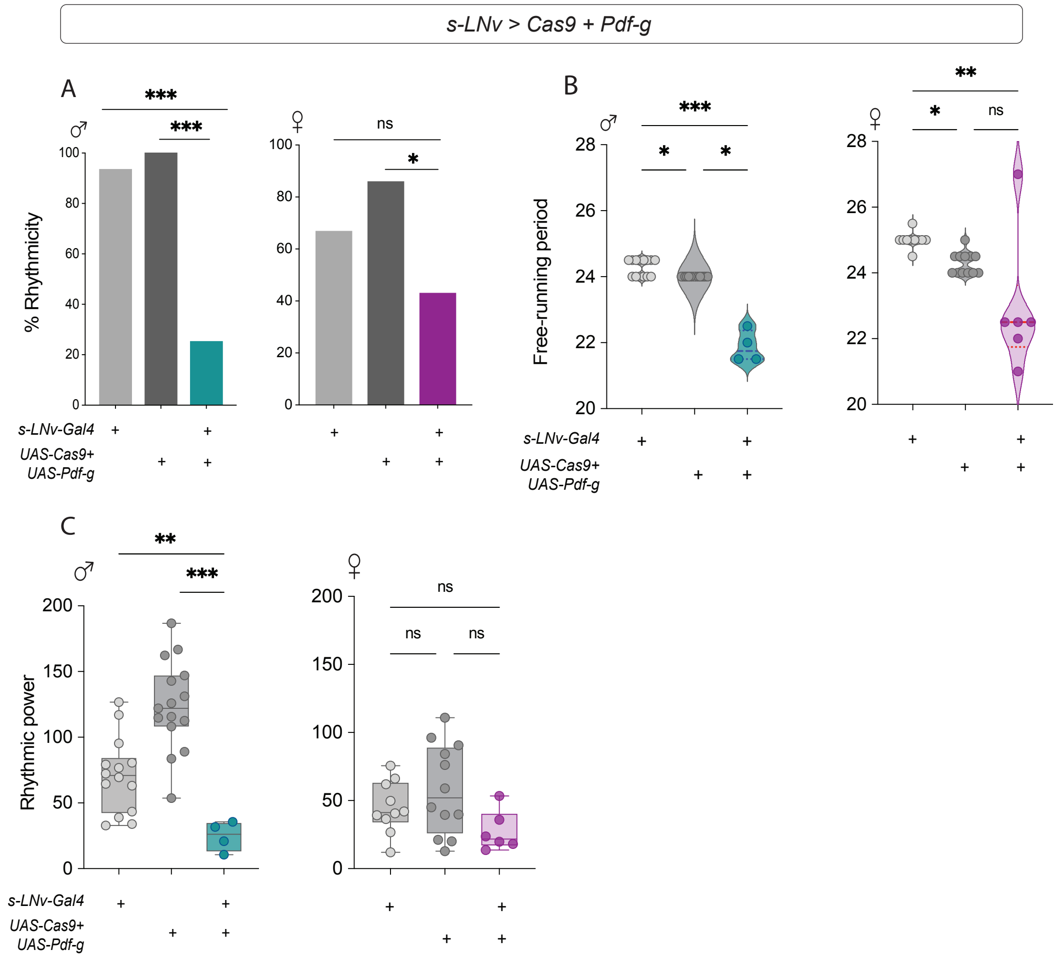

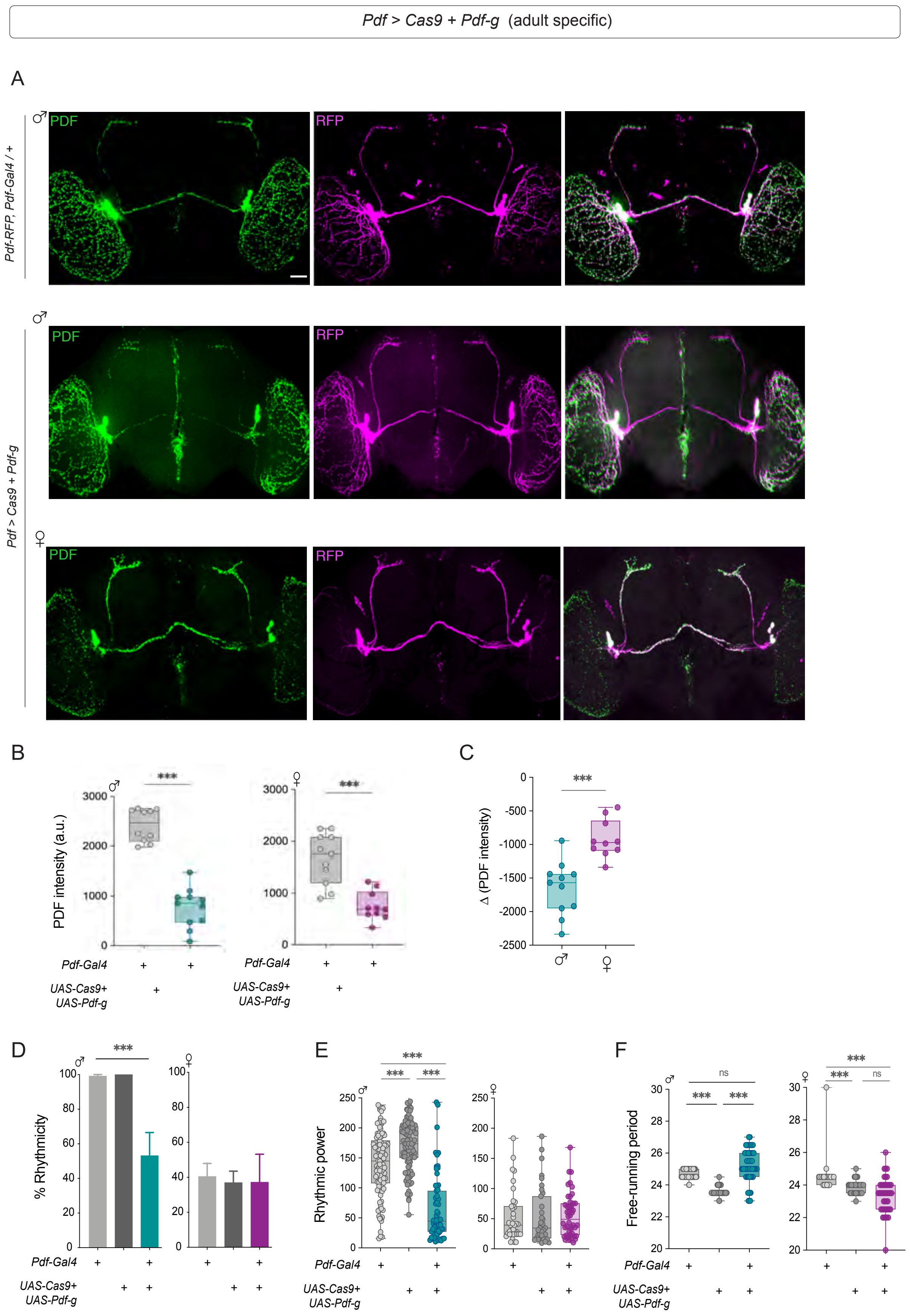

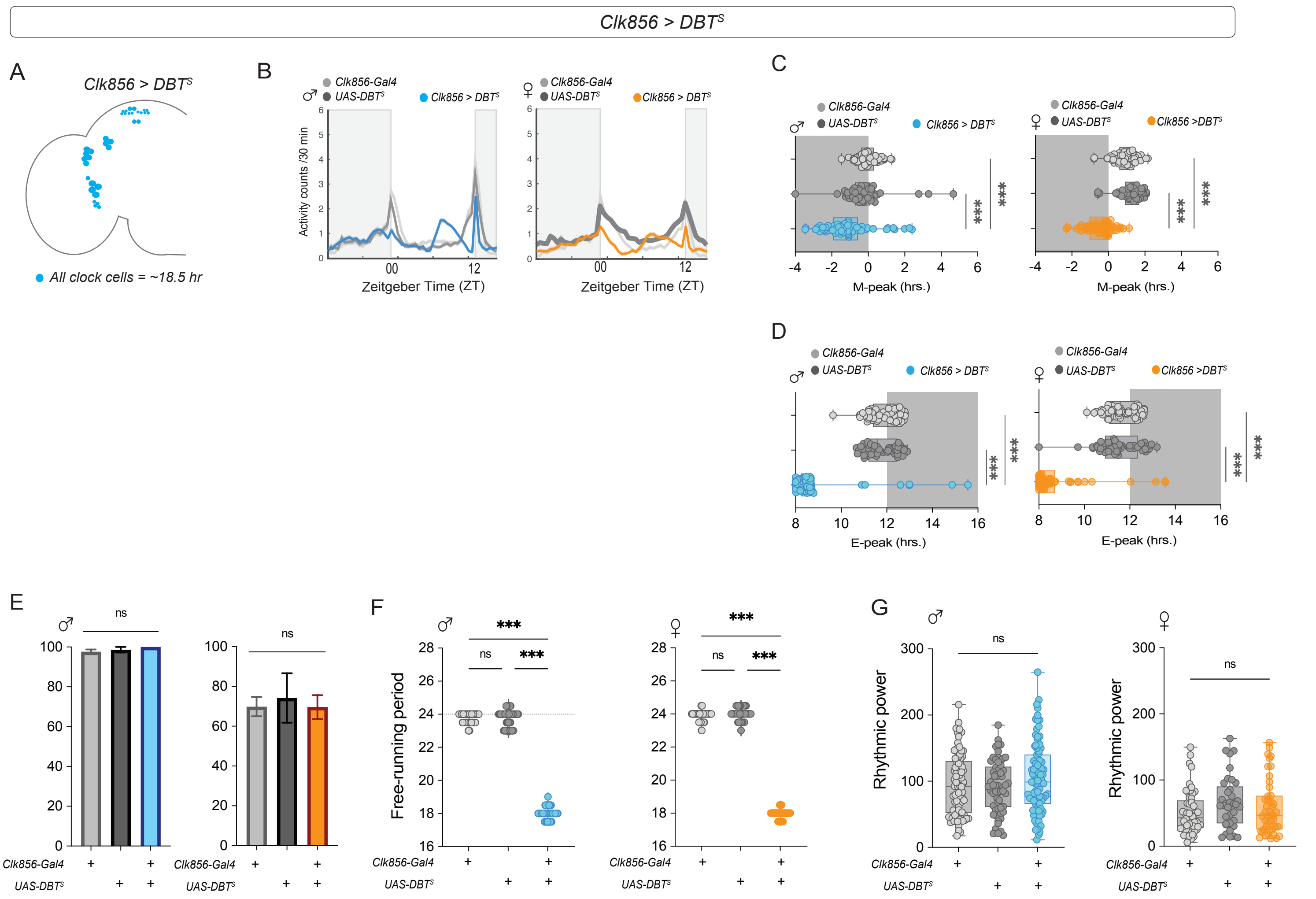

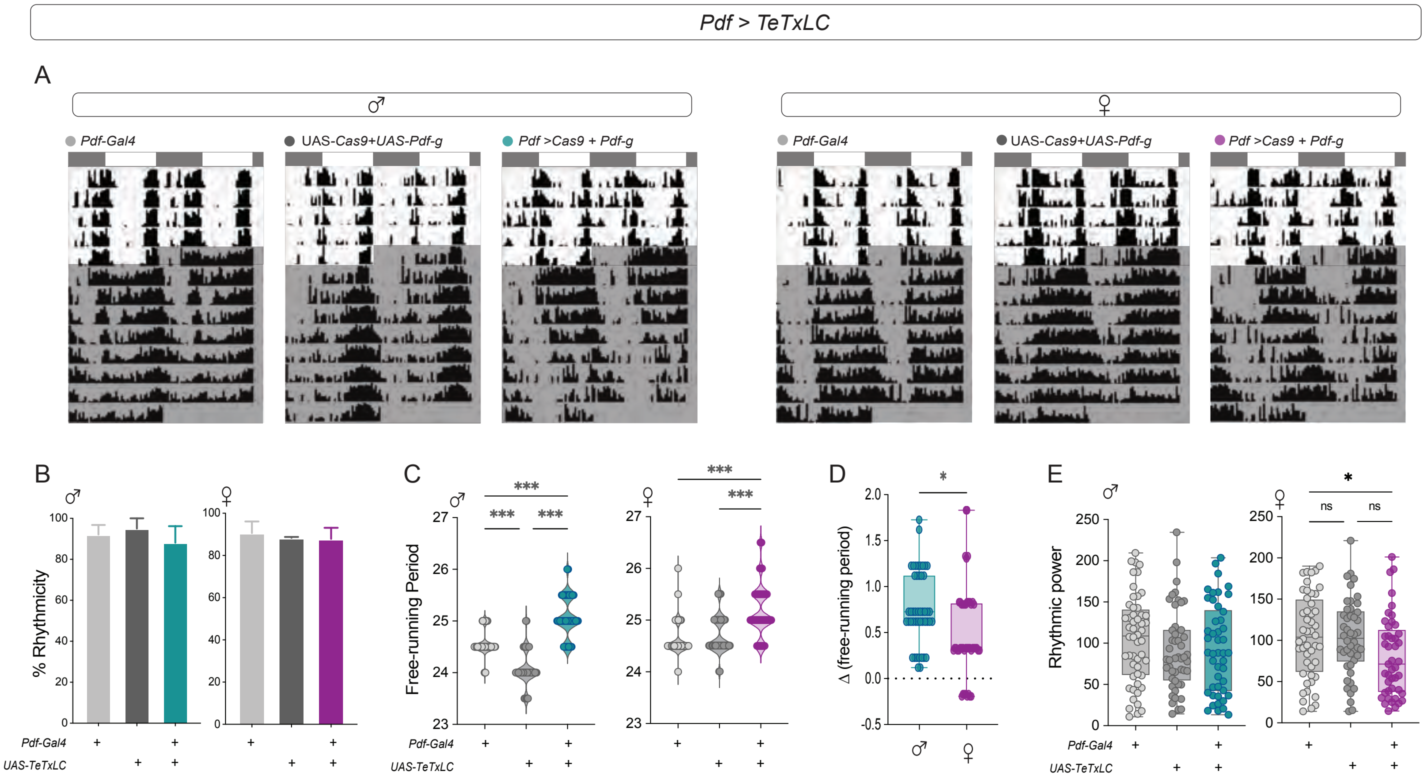
